# Do NSm virulence factors in the Bunyavirales viral order originate from Gn gene duplication?

**DOI:** 10.1101/2023.04.13.536692

**Authors:** Victor Lefebvre, Ravy Leon Foun Lin, Laura Cole, François-Loïc Cosset, Marie-Laure Fogeron, Anja Böckmann

**Author notes:** Correspondence (Fogeron, M.-L.); (Böckmann, A.).

## Abstract

One-third of the nine WHO shortlisted pathogens prioritized for research and development in public health emergencies belong to the *Bunyavirales* order. Several *Bunyavirales* species carry an NSm protein that acts as a virulence factor. We predicted the structures of these NSm protein and unexpectedly found that in two families, its cytosolic domain is inferred to have a similar fold to the cytosolic domain of the viral envelope-forming glycoprotein N (Gn^cyto^) encoded on the same genome fragment. We show that although the sequence identity between the NSm^cyto^ and Gn^cyto^ domains is low, the conservation of the two zinc finger-forming CCHC motifs explains the predicted structural conservation. Importantly, our predictions provide a first glimpse into the long unknown structure of NSm and its link to virulence. Also, these predictions suggest that NSm is the result of a gene duplication event in the *Bunyavirales Nairoviridae* and *Peribunyaviridae* families, and that such events may be common in the recent evolutionary history of RNA viruses.

## Introduction

Viral research is often conducted in a situation of urgency, going from knowing virtually nothing about a virus to understanding it as quickly as possible. To change this inefficient situation, the World Health Organization (WHO) has decided on a list of ten pathogens for which research should be prioritized. These pathogens have a high pandemic potential due to their lethality and the lack of effective treatment or prevention. They are mainly transmitted by vectors such as mosquitoes and ticks, with which humans will increasingly interact in the current context of climate change. *Bunyavirales* represent thus a persistent global health challenge, and research on these pathogens is important to prevent future outbreaks or even an increase in their spread, which could have devastating consequences. The *Bunyavirales*, a large viral order, provide three of these pathogens (Lassa (LAS), Rift Valley

Fever (RVF), and Crimean-Congo Hemorrhagic Fever (CCHF) viruses). Although research is now intensifying at all levels, it still lags far behind that on other shortlisted viruses. A virulence factor present in many pathogens of the order, the non-structural protein m (NSm), has roles in antagonizing immune responses, promoting viral assembly and infectivity, and maintaining infection in their transmission vectors [1]. Although a pathogenic virus in the order may not have such an NSm protein and a virus containing an NSm is not necessarily a pathogen, NSm has been shown to play a clear role as an amplifier of infection in some of the most threatening viruses, such as CCHFV. Such proteins are thus prime drug targets and attenuated viruses carrying deletions of these proteins can serve as a basis for vaccine development. Knowledge of the structure of the NSm protein is therefore central to understand disease progression and viral pathogenesis, and to develop strategies for intervention and treatment.

The viruses of the order recognize a wide variety of hosts, including humans, in which they can cause severe diseases. Arthropod vectors such as ticks, mosquitoes and sand flies present the most common modes of transmission, which can also be direct, by contact with infectious blood or body fluids (Figure 1a). The *Bunyavirales* are diverse but share a single-stranded negative-polarity RNA genome, often with the three segments L, M and S (Figure 1b). They encode at least four structural proteins that make up the viral particle: the envelope glycoproteins Gn and Gc; the NP nucleoprotein, and the L viral RNA-dependent RNA polymerase. The structural and functional similarities in bunyaviruses have been reviewed recently, in order to evaluate perspectives for pan-bunya antivirals [2].

**Figure 1.**
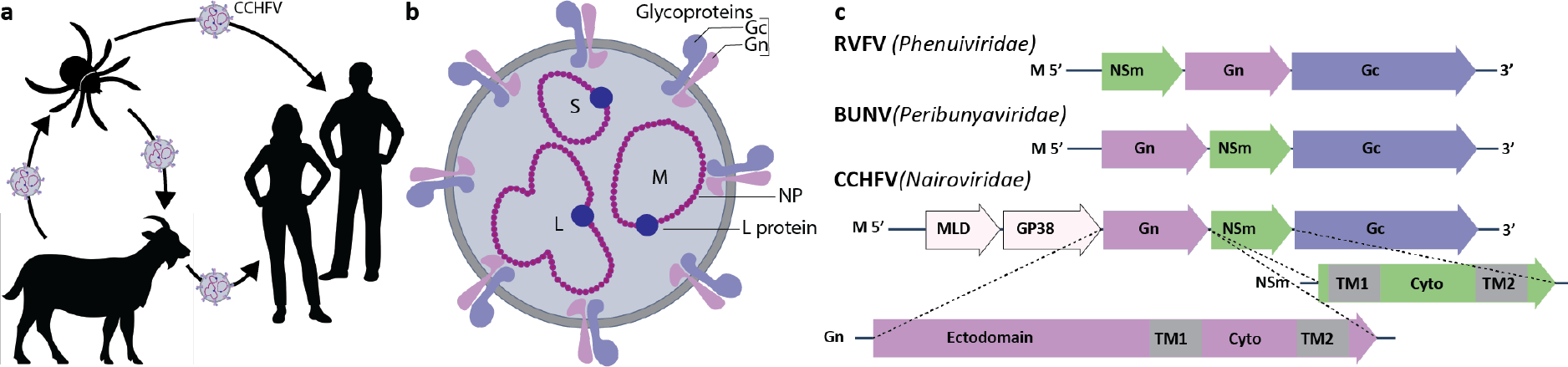
The *Bunyavirales*. a) Transmission modes at the example of CCHFV, where the virus transits between the tick vector, life stock and humans. b) The CCHFV viral particle. c) The NSm non-structural protein is encoded on the M gene of certain Bunyavirales species [3].

NSm is, *ex vivo*, an accessory protein [1] encoded on the M segment in the *Nairoviridae, Peribunyaviridae*, and *Phenuiviridae* families of the order [3]. For CCHFV (*Nairoviridae* family) and Bunyamvwera virus (BUNV, *Peribunyaviridae* family), NSm is the C-terminal cleavage product of Pre-Gn [4,5] (Figure 1c). In the *Phenuiviridae* family, as in RVFV [1], NSm precedes Gn. NSm has 150-200 residues, has been described to be twice membrane-spanning, and localizes mainly to the Golgi in infected cells [6]. The structure of NSm has long remained a mystery. Today, as we show here, artificial intelligence structure prediction programs such as AlphaFold2 [7] can provide a first insight.

## Materials and Methods

An implementation of AlphaFold2.1 [7] software on a local server was used for predictions. Amino-acid sequences were entered and predictions were run in the monomer mode. Amino-acid sequences were obtained from the UniProt data base. Resulting AlphaFold archives are provided upon request. Per-residue estimates of the confidence of the models are given on a scale from 0-100 in the individual PDB files of the predictions (pLDDT). Color coding in the Figures goes from darkblue (100), light blue (90), yellow (70), orange (50), to red (0). pLDDT > 90 indicate a model with high accuracy; 70 and 90 a generally good backbone prediction; 50 and 70 low confidence, and < 50 not reliable. pLDDT values for the different models are summarized in Table S1. Predicted structures were visualized, and also matched where applicable, using ChimeraX-1.5 [8], which was also used to calculate backbone RMSDs. Sequence alignments were done using the multiple protein sequence alignment tools available through the NPS@ web interface [9].

## Results

### NSm resembles the cytosolic domain of glycoprotein Gn in the Nairo- and Peribunyaviridae

Figure 2a shows an AlphaFold2 prediction of CCHFV NSm. The resulting model shows two transmembrane domains and one globular domain. As can be seen from the color coding, the cytosolic domain shows high accuracy, while the prediction of the transmembrane domains shows lower confidence. The cytosolic domain surprisingly shows a very well-defined fold, predicted with high confidence. A closer look at the AlphaFold2 hit report interestingly shows that the previously determined CCHFV Gn cytosolic domain (Gn^cyto^) structure (PDB 2L7X [10]) represents the best hit in hhsearch (as provided in the AlphaFold2 result files). A comparison of CCHFV NSm and Gn^cyto^ (in magenta in Figure 2a) shows that Gn^cyto^ appears to be indeed fully reproduced in the predicted NSm model (backbone RMSD of 1.6 Å). An alignment of the CCHFV Gn and NSm cytosolic domain sequences from different strains (Figure 2b and Table S2) reveals the rationale behind the high similarity: while less than 22 % of the sequence is identical, the CCHC motifs of the two ββα zinc fingers (ZFs) in Gn^cyto^ are indeed fully conserved between the two proteins, as also shown in the zoom in Figure 2c. Comparing CCHFV NSm with another member of the *Nairoviridae* family, Figure 2d shows the alignment (see also Table S3) and AlphaFold2 models of CCHFV and Dugbe virus (DUGV) NSm^cyto^, which overlap nearly perfectly (RMSD of 1.8 Å).

**Figure 2.**
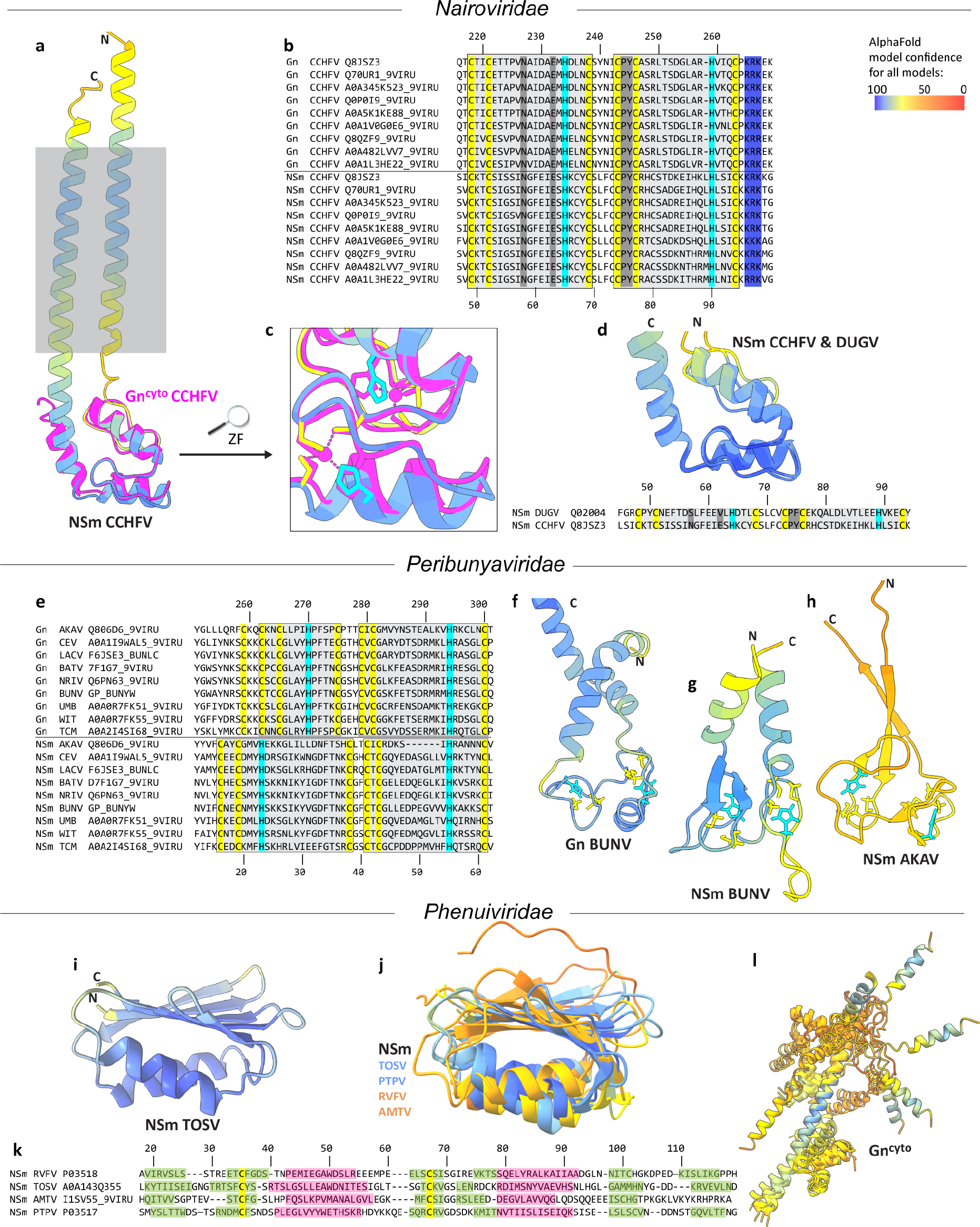
AlphaFold2 structure predictions and sequence alignments of the *Bunyavirales* NSm and Gn^cyto^ proteins. a) CCHFV NSm model overlayed with Gn^cyto^ (PDB 2L7X [10]) in magenta, with the grey box indicating the lipid bilayer. Colors reflect the reliability of the prediction as assessed within AlphaFold pLDDT values, from blue (reliable) to red (poor), as shown in the bar at the top right. b) Sequence alignments of Gn^cyto^ and NSm^cyto^ of different CCHFV strains (for the full alignment and numbering of the M polyproteins see Table S2). Residue numbering is given for Q8JSZ3. Conserved residues are highlighted: yellow, Cys; cyan, His; blue, basic residues; grey, other. c) The double ZF motif conserved in NSm and Gn. d) Sequence alignments and AlphaFold2 models of DUGV and CCHFV NSm (for full CCHFV/DUGV M polyprotein alignment see Table S3). e) Gn and NSm sequence alignment for the *Peribunyaviridae* (for the full alignment and numbering of the M polyproteins see Table S4). AKAV, Akabane virus; CEV, California Encephalitis Virus; LACV, La Crosse Encephalitis Virus; BATV, Batai Virus; BUNV, Bunyamwera virus; NRIV, Ngari Virus; UMBV, Umbre Virus; WITV, Witwatersrand Virus; TCMV, Tacaiuma Virus. Structure predictions (f, g) reveal less similarity of Gn^cyto^ and NSm^cyto^ than in the *Nairoviridae*. AKAV NSm (h) shows poor prediction accuracy. i) *Phenuiviridae* NSm is predicted in a highly accurate fold for TOSV and PTPV, but less so for RVFV and AMTV (j). k) Manual sequence alignment of the NSm domains of different members of the *Phenuiviridae family* taking into account predicted secondary structures: α-helices, pink; β-strands, green. Conserved Cys are highlighted in yellow. l) Gn^cyto^ model showing poor accuracy and poor similarity to NSm. Colors in all models give AlphaFold2 pLDDT values, from blue (reliable) to red (poor), as shown in the bar at the top.

The *Peribunyaviridae* family also carries NSm in its M gene (Figure 1c). A sequence alignment (Figure 2e; for full alignment see Table S4) here reveals the full conservation of seven cysteines and two histidines in Gn^cyto^, and six cysteines and two histidines in NSm^cyto^. Sequence identity between Gn and NSm is with 19 % similar to the *Nairoviridae*. In contrast to the *Nairoviridae*, there is no experimental structure for *Peribunyaviridae* Gn^cyto^. The pattern that can be postulated in Gn^cyto^ as a C-terminal zinc finger (ZF2, right grey box) is mainly conserved in NSm, resulting in a possible CCHC motif. The situation is less obvious for the pattern corresponding to a putative N-terminal zinc finger (ZF1), which in Gn^cyto^ could be either Cys3-His, Cys2-His-Cys, or even Cys4, while it must be Cys2-His-Cys in NSm. The predicted AlphaFold2 models for BUNV Gn and NSm are shown in Figure 2f and g. A double zinc finger is predicted for both Gn^cyto^ and NSm, with two CCHC motifs. ZF2 in Gn^cyto^ shows a ββα fold like CCHFV Gn^cyto^, while ZF1 is similar to that in HIV-1 NCp7 [11], which also shows a Cx2Cx4Hx4C sequence. The RMSD between the two models is 5.8 Å. The resulting models of the other aligned family members are very similar (not shown), with the exception of AKAV NSm, which presents a very different fold, probably due to the six-residue deletion in ZF2, and is predicted with low confidence (Figure 2h). Thus, in contrast to the *Nairoviridae*, the predicted structures for Gn^cyto^ and NSm^cyto^ seem to differ at least partially from each other. The Gn^cyto^ ZF2 ββα fold is surprisingly not formed in NSm, despite the very similar length of the loop between the CC and HC parts of the motif. ZF1 is completely different, as already suggested by the different sequence motifs. However, it must be remembered that, in contrast to the *Nairoviridae*, where the Gn^cyto^ structure is known [10], there is no experimental structure for either protein that could assist AlphaFold2 predictions in this family. Thus, to obtain experimental data will be decisive in further analyses.

### NSm models in Phenuiviridae do not resemble Gn

The third family carrying an NSm protein is the *Phenuiviridae* family. The NSm sequences do not suggest the presence of a zinc finger, as only two conserved cysteine residues, and no histidine, are present in RVFV, Toscana virus (TOSV), Arumowot virus (AMTV) and Punta Toro virus (PTPV) NSm (Table S5,6). The identity in the alignment between the different NSm is less than 3 %. AlphaFold2 predictions however surprisingly lead to similar structural models of the NSm proteins, as shown in Figure 2i,j, where TOSV NSm is shown in isolation and then superimposed on the three other structural models. In addition to TOSV, the structure of PTPV is predicted quite reliably, whereas RVFV and AMTV are judged to be less accurate. The resulting folds include two α-helices and a six-stranded β-sheet. The pairwise RMSD for the proteins is 2.3 Å (PTPV to TOSV), 4.4 Å (RVFV to TOSV), and 10 .0 Å (AMTV to TOSV). Accordingly, Figure 2k shows a manual alignment of the protein sequences taking into account the secondary structural features of the predicted models. The two cysteines remain approximately aligned, but do not form a disulfide bond with each other. While the folds look similar, the fact that there is no experimental structure across the different NSm in the *Phenuiviridae* also seems to result in the lack of a clear model to use in the predictions. Experimental data will be crucial here as well.

Gn^cyto^ in the *Phenuiviridae* is predicted as multiple α-helices, forming a not very well-defined 3D fold (Figure 2l). Identity within the different Gn^cyto^ is less than 5 % (Table S7). While the orientation of the helices with respect to each other is not maintained in the different proteins, the α-helical structures are rather consistent. Still, one can see that the reliability in general is poor, as indicated by the color coding, besides the light blue helical fragment shown in the center, and on which the structures were matched. The predicted models clearly indicate that there is no relationship between NSm and Gn^cyto^, as also reflected in the alignment, which shows less than 10 % identity (Table S8).

## Discussion

The structure predictions of the different cytosolic domains of Gn and NSm reveal a common fold in two out of the three families investigated: the *Nairoviridae* and the *Peribunyaviridae*. This observation correlates with the position of NSm on the genome, which follows Gn in these two families, but precedes Gn in the third one, the *Phenuiviriade* family (Figure 1c). It is unclear how NSm originated; the predicted structural similarity, which is most striking in the *Nairoviridae*, leads us to hypothesize that NSm may have originated there from a duplication event of a gene fragment from Gn^cyto^. Gene duplication is a process by which a genetic sequence, or a fragment thereof, is copied, creating an additional gene sequence [12]. This can occur naturally through mutations in DNA replication or through recombination. Both the mechanisms that can lead to the formation of gene duplicates, as well as the fates of the new duplicated genes are wide-ranging and can depend on several factors [13]. Duplication of genes or gene fragments can lead to the evolution of new functions of the duplicated fragment or to an increase in the amount of protein produced by it [12]; both can result in increased virulence. In viruses, the frequency of gene duplication can vary greatly depending on the specific virus and the conditions under which it replicates [14]. However, duplication of gene fragments is common in many viruses. In particular, RNA viruses can undergo frequent duplication of gene fragments due to their rapid replication cycles, recombinational potential and error-prone replication mechanisms. And indeed, previously, a duplication event of a gene fragment from Gn was proposed to have resulted in CCHFV in the virulence factor GP38 [15] and an internal gene fragment duplication has been described within the CCHFV Gc [16]; for both GP38 and Gc, experimental 3D structures are described. Gene duplication in viruses remains indeed difficult to nail down based solely on sequence information. We here show that structural knowledge, including from predictions, can be decisive to identify possible duplication events.

One could mention that another possible hypothesis would be that gene duplication started from NSm, and not Gn. This is however unlikely since Gn is a structural, and thus necessary protein; also, there are viruses in the *Bunyavirales* for which no NSm was identified, such as the *Hantaviridae*, where Gn^cyto^ however shows a highly similar double-zinc-finger motif as in CCHFV [17,18]. Convergent evolution from a host protein could also be considered as possible origin; however, as already discussed by Estrada and coworkers [17,18], the Gn, and thus also the NSm, double zinc-fingers do not fold independently as classical zinc fingers do, but each finger affects the folding of the other [17,18]. Few intimately contacting double zinc fingers have been reported in the PDB, one example being the yeast Bcd1 protein [19], which however shows a different 3D fold than CCHFV Gn^cyto^.

For the *Phenuiviridae*, our models suggest that Gn gene fragment duplication is not the origin of NSm, and the protein, with a completely different structural organization than Gn^cyto^, must have been acquired by another mechanism, possibly from the vector host, considering its possible role in maintaining infection in viral vectors [20].

The Gn^cyto^ zinc fingers are believed to be involved in nucleic acid interactions with the viral genome, or also the nucleoproteins [10,17,18]. The exact role of NSm in CCHFV infection is still unclear, and whether NSm has evolved for new purposes or rather for supporting certain functions of Gn by increasing their level remains to be determined. One could mention that zinc fingers have been shown to play an important role in the development of hepatocellular carcinoma (HCC) [21]. Indeed, CCHFV can cause severe liver injury in humans [22,23]; whether NSm and/or Gn^cyto^ zinc fingers play a role in liver injury in CCHFV infection remains to be elucidated.

## Conclusions

Based on current knowledge of the phylogenetic tree of evolutionary relationships among different viral lineages of the order Bunyavirales [1], it is unclear how NSm originated and evolved, making it difficult to trace its origin. Our analysis, based on structure prediction, suggests that gene fragment duplication events in the *Nairoviridae* and *Peribunyaviridae* families might have led to NSm. Experimental structural data will however be needed to confirm this hypothesis. With gene duplication already reported for GP38 and Gc in CCHFV, our work adds NSm as a candidate product from such an event. While duplication of gene fragments has rarely been reported in RNA viruses for proteins where no 3D structures are available, this may be due to the high mutation rates in these viruses; as a consequence, the resulting low protein sequence similarity can significantly complicate a conservative sequence search approach. Thus, the present analysis not only provides first structural models for the enigmatic NSm proteins, but also highlights the importance of structure predictions to identify duplications of gene fragments in RNA viruses.

## Supplementary Materials

Table S1: pLDDT values for the different Alphafold predictions

Table S2: Sequence alignment of the M polyprotein of different CCHFV strains

Table S3: Sequence alignments and numbering of CCHFV and DUGV M polyproteins

Table S4: Sequence alignments and numbering of different *Peribunyaviridae* M polyproteins

Table S5: Sequence alignments of different *Phenuiviridae* M polyproteins.

Table S6: Sequence alignment of *Phenuiviridae* NSm

Table S7: Sequence alignment of *Phenuiviridae* Gn^cyto^

Table S8: Sequence alignment of RVFV Gn^cyto^ and NSm

## Author contributions

VL, RLFL and LC performed sequence analyses; VL, RLFL performed bioinformatics analyses and alignments; VL and AB did structure predictions; FLC advised on CCHFV virology; MLF and AB designed the research. AB and MLF wrote the paper, with input from all authors.

## Acknowledgement

This work was supported by the Agence Nationale de la Recherche (ANRS ECTZ 205074, ECTZ 158948), by the RESPOND program of the LabEx Ecofect (ANR-11-LABX-0048) of the “Université de Lyon”, within the program ‘‘Investissements d’Avenir” (ANR-11-IDEX-0007) operated by the French National Research Agency (ANR) and by an API-2022 MMSB.

## Data Availability Statement

All AlphaFold models are available from the corresponding authors on request.

## Conflict of interests

The authors declare no conflict of interest.

## Notes

### Competing Interest Statement

The authors have declared no competing interest.

### Summary of Updates

-consistent AlphaFold pLDDT coloring -contains a more homogenous description of the structures and alignments -adds sequence alignments in the supplementary materials

