## Supporting Information for "Do NSm virulence factors in the Bunyavirales viral order originate from Gn gene duplication?"

**Table S1: pLDDT values for the different Alphafold predictions**

| Protein | Virus | pLDDT of relaxed models |
|---|---|---|
| *Nairoviridae* | | |
| $Gn^{cyto}$ | CCHFV | "model_1": 86.09535594068562, "model_2": 84.13366683662879, "model_3": 83.88211244700702, "model_4": 83.05718111553344, "model_5": 85.9895667335572 |
| NSm | CCHFV | "model_1": 73.20299605327742, "model_2": 76.49797744544017, "model_3": 75.93087992726971, "model_4": 76.0229989062226, "model_5": 65.98470683947802 |
| NSm | DUGV | "model_1": 70.79763901984666, "model_2": 70.76243810168731, "model_3": 75.502799092311, "model_4": 67.2813735126672, "model_5": 72.25758629063004 |
| *Peribunyaviridae* | | |
| $Gn^{cyto}$ | AKAV | "model_1": 87.1395655314573, "model_2": 85.33816484988238, "model_3": 84.7078396197587, "model_4": 86.09484027753327, "model_5": 87.35710551505787 |
| $Gn^{cyto}$ | LACV | "model_1": 81.77028814252233, "model_2": 83.44042634487546, "model_3": 82.36054330594433, "model_4": 84.90695096535228, "model_5": 85.64488588007816 |
| $Gn^{cyto}$ | BUNV | "model_1": 85.12077569987977, "model_2": 82.90528866481296, "model_3": 86.44181082140551, "model_4": 86.22130146282308, "model_5": 86.18086927041759 |

| | | |
|---|---|---|
| Gn$^{cyto}$ | TCM | "model_1": 85.58792842770676, "model_2": 84.16999856616901, "model_3": 88.36289501162956, "model_4": 87.34276619491422, "model_5": 88.34218834238854 |
| NSm | AKAV | "model_1": 76.23471541210738, "model_2": 70.51273047640764, "model_3": 73.76506337951135, "model_4": 74.04727772569176, "model_5": 70.6300246037283 |
| NSm | CEV | "model_1": 78.6825363461259, "model_2": 74.57560897328426, "model_3": 80.97233719972321, "model_4": 73.39347604833836, "model_5": 70.89139020241463 |
| NSm | LACV | "model_1": 84.40982502730394, "model_2": 84.64757333337988, "model_3": 85.63314774917781, "model_4": 84.26046572656772, "model_5": 75.42250934288181 |
| NSm | BATV | "model_1": 88.99156581198358, "model_2": 88.36044208391598, "model_3": 89.55390238139809, "model_4": 88.58448280826944, "model_5": 88.71501308857546 |
| NSm | NRIV | "model_1": 91.09281025506803, "model_2": 90.40338236338134, "model_3": 91.1631716079305, "model_4": 90.64013308258669, |
| NSm | BUNV | "model_1": 82.10726193774362, "model_2": 78.38550639506562, "model_3": 77.08859780341184, "model_4": 78.75972713981923, |

| | | |
|---|---|---|
| | | "model_5": 74.65827632820661 |
| NSm | UMB | "model_1": 77.93705087566654, "model_2": 66.43847386407046, "model_3": 72.63109678782779, "model_4": 76.95066852603681, "model_5": 67.99364777647244 |
| NSm | WIT | "model_1": 76.92499302901366, "model_2": 77.59385380361836, "model_3": 81.30708582371636, "model_4": 77.17394526060112, "model_5": 78.87536381348019 |
| NSm | TCM | "model_1": 85.3635576239163, "model_2": 86.62886020335853, "model_3": 89.55647536875307, "model_4": 87.29175285543657, "model_5": 82.79121744782258 |
| *Phenuiviridae* | | |
| $Gn^{cyto}$ | RVFV | "model_1": 69.61977603515074, "model_2": 68.85799880213563, "model_3": 70.23997304046667, "model_4": 70.20191467985602, "model_5": 67.947864220289 |
| $Gn^{cyto}$ | TOSV | "model_1": 65.85872776445585, "model_2": 64.24228565182545, "model_3": 70.11255025211632, "model_4": 65.93771674937094, "model_5": 62.91438040700215 |
| $Gn^{cyto}$ | AMTV | "model_1": 58.38951112814995, "model_2": 53.678962596338806, "model_3": 54.925130043034024, "model_4": 54.94835536789018, "model_5": 50.896421376769645 |
| $Gn^{cyto}$ | PTPV | "model_1": 72.30293286991761, |

| | | |
|---|---|---|
| | | "model_2": 70.0753852989372, "model_3": 72.71336378248058, "model_4": 73.97087128823867, "model_5": 68.94298121981585 |
| NSm | RVFV | "model_1": 48.76205757798705, "model_2": 44.78753435573813, "model_3": 49.310055039113905, "model_4": 42.37059916214862, "model_5": 29.711595969397308 |
| NSm | TOSV | "model_1": 73.69630562899772, "model_2": 65.60234422942727, "model_3": 76.42367296131455, "model_4": 61.62299240563712, "model_5": 73.19451202003762 |
| NSm | AMTV | "model_1": 41.26781514681544, "model_2": 32.44954707495688, "model_3": 40.45080997902588, "model_4": 33.96799675844322, "model_5": 36.48935377673723 |
| NSm | PTPV | "model_1": 77.93808623801871, "model_2": 77.57371349467493, "model_3": 78.02042218764377, "model_4": 77.58799828079687, "model_5": 77.55586181363482 |

**Table S2**: **Sequence alignment of the M polyprotein of different CCHFV strains**. The $Gn^{cyto}$ and NSm zinc finger domains are highlighted in green and yellow, respectively, within $Gn^{cyto}$ and full-length NSm in grey. Red type/ asterisk (*), green type/colon (:), and blue type/dot (.) indicate identical amino acid residues, conserved substitution, and semi-conserved substitutions respectively.

```
                            10        20        30        40        50        60        70        80        90       100
                             |         |         |         |         |         |         |         |         |         |
Q8JSZ3              -----MHISLMYAILCLQLCGLGETHGS----HNETRHNKTDTMTTPGDN-PSSEPPVSTALSITLDPSTVTPTTPASGLEGSGEVYTSPPITTGSLPL-
Q70UR1_9VIRU        -----MHISLMYAVFCLQLCGLGKTNGP----HNGTEHNNTHVMTTPDDS-QSPEPPVSTALPVTPDPSTVTPSTPASGLEGSGEVYTSSPITTKGLSL-
A0A345K523_9VIRU    MPINIMHTLLVCFILYLQLLGPGGAHGQ----SNATEHNGTNTTTAPGTS-QSPKPPASTTPSHAPEPSTIKLTTPTSETEGSGET-TPPTNTTQDSPP-
Q0P0I9_9VIRU        MPTNIMHTLLVCFILYLQLLCLGGAHGQ----LNTTEHNGTNNTTAPGAS-QSPKPPMSTTPPHAPESSTIKPTTPISEAEGSGET-TSTPNTTQGLSS-
A0A5K1KE88_9VIRU    MPISIMHISLMCAVLCLQLYGLGGTHGS----HNGTEHNKTDAVTTSSDS-QSSEPPVRTTQPVTPDSSTVTPTTPASVPEGSGEVYTSPLNTTEGPSP-
A0A1V0G0E6_9VIRU    MLRYILYALLASAILHQHLYKVGADTQKPTVRTNTT-HKISPTATANRTTYEQATPSTTPKSSTKP--TTHIATAHLAESDGSGETTLLLPTTASQYTT-
Q8QZF9_9VIRU        MLFHFKFMLLINFILCHLLWGGGGVTGGVETNSSSTQAIQTPPVTSNSTT---PGPSTDTATETASSMTTSTPDTQVTTDNGSGEPSTDPLTTNEATPPL
A0A482LVV7_9VIRU    MLFHFKFMLLINFILCHLLWGGVGVTGGVETNSSSTQAIPTPPVTSNSTT---PGPSTDTATETASSMTTSTPDTQVTTDNGSGEPSTDPLTTNEATPPL
A0A1L3HE22_9VIRU    -MFCLKLVLLINLTLCQLLEGNDRVTSADGINSNTTQTNLGISTILNSTT-PAPGPSTDTTMETVPSTADSMLETQVTTDSGSGEPSTEPTTTGEATSST
                             *    :   *              . *             .   . *.  .        :     :  :  .****       *    .
Prim.cons.          M2F2IMHILL222ILCLQL4GLGG22G3VETNSNTTEHN2TPT2T22ST2YQSP2PPTST233TAP2PST2TPTTP2S2DEGSGE32TSPPTTTE222PL

                           110       120       130       140       150       160       170       180       190       200
                             |         |         |         |         |         |         |         |         |         |
Q8JSZ3              -----------SETTPELPVTTGTDTLSAGDVDPSTQTAGGTSAPTVRTSLPNSPSTPSTPQDTHHPVRNLLSVTSPGPDETSTPSG-TGKESSATSSPH
Q70UR1_9VIRU        -----------PEATSEPPATTSVVTSSASDTDSSTQAAGDTPTPTVRTSLPSSPSTPSTSQGTHYPVRSLLSVTSPKPEETPTPSK-SGKDDLATNSPH
A0A345K523_9VIRU    -----------LETTPERSATTATSTSGTDNMNSTTQTTDNTPTSTVSTSLSSSPSTPSTSQGMHHPVRSLLSVSSSMTATTPTPMS-PGEMSSETSSQH
Q0P0I9_9VIRU        -----------PETTSERPATTAISTSSTDSTNPTTQMTDNTPTLTVSTSPSSSPSTPSTPQGIYHPARSLLSVSSPKTVTTPTPTS-PGEMSSETSSQH
A0A5K1KE88_9VIRU    -----------PESTPESPVAASTGTPSASDVNSSTQAARDTSTPTVRTSPPNSLSTPPTPQETHHPVRNLLSVTSHKPEETTAPSE-SGKESPAANSPH
A0A1V0G0E6_9VIRU    ----------EGQTSLEPTAEETGSNTIPSDTTSTSPITENSESTDTQMNVSDSSSSPLATQGARSPARNLLSVPSVEPGQQTTPKQPTTKGTVTPTTPQ
Q8QZF9_9VIRU        SVTTAQTTETKKHTTLDPSTSTNPDATTPSITILSASNTSLPISS-VHASPSPSSSEEPTPQNAHHLSRSLLSAVSTETGQIASAAT-AAKEITTKNSPP
A0A482LVV7_9VIRU    SVTTAQTKETKKHTTLDPNTSANPDATTPSITILSASNTSLPTSS-VHASPSPSSSEEPTPQNAHHLSRSLLSAVSTETGQTASATT-ATKEITTKNSPP
A0A1L3HE22_9VIRU    TAATVSMQETKKHMTQNTRSTADPDTTTPNSTTSVEPTTIPPTSPIPQATPTPGHGKAPTLPNTHHLSRSLLSAVSTEAGQTTPATS-TTEAASAPSSSS
                                . : :           ..        :  . :     . . . .   :        *.***. *  .   ...   . :     .:
Prim.cons.          SVTTAQT3ETK2ETT2EPPATT32DTTSPSDTNSSTQTT3NTP2PTVRTSPS2SPSTP2TPQGTHHPVRSLLSV2S2E2GQT2TP2SPTGKESSAT2SPH

                           210       220       230       240       250       260       270       280       290       300
                             |         |         |         |         |         |         |         |         |         |
Q8JSZ3              PVSNRPPTPPATAQGPTENDSHNATEHPESLTQSATPGLMTSPTQIVHPQSATPITVQDTHPSPTNRSKRNLKMEIILTLSQGLKKYYGKILRLLQLTLE
Q70UR1_9VIRU        PATSRPTTPPTTAQKPTENNSHNTTEQLESLTHLATLGLMISPTQTVLPQSVTSIAIQDIHNSPTNRSKRNLDMEIILTLSQGLKKYYGKILKLLHLTLE
A0A345K523_9VIRU    PAMSRTPTPHTATQVSTENANRSTSRQSESSAQPATQSPVTSPVQSILLMSATPTTIQNIHPSPTNRSKRNLEMEIILTLSQGLKKYYGKILKLLHLTLE
Q0P0I9_9VIRU        SAMSRIPTPHTATRVSTENTNHSTPRQSESSAQQTTPSPMTSPAQSILLMSAAPTAVQDIHPSPTNRSKRNLETEIILTLSQGLKKYYGKILKLLHLTLE
A0A5K1KE88_9VIRU    PVASRPPTPSTTAQGPTENNSHSTTGQLESLTQSTTLDPMTSLTQTVLPQSVTSIAVQDTHPSPTNRFKRNLEMEIILTLSQGLKKYYGKVLRLLQLTLE
A0A1V0G0E6_9VIRU    PTTSSPTPAPTSATLSTNSSSISTPVQTSSPLQPMAPSHETQHALYTTSTESTLRTLSAERIHPTNRSRRNSEVEIILTLPQGLKKYYSRLLKLLHLTME
Q8QZF9_9VIRU        TAGND-----TMVTSAPNGNIRNTPEQANPTAPEASPKIQTTQNQQTVTPEATPITTITALTDLTNRSKRDSKVQIILTFSQGLKKYYSKILKLLQLTQE
A0A482LVV7_9VIRU    TAGND-----TMVTSAPNGNTRNTPEQANPTAPEASLKIQTTQNQQTVTPEATPITTVTALTDLTNRSKRDSKVQIILTFSQGLKKYYSKILKLLQLTQE
A0A1L3HE22_9VIRU    TAENN-----TVSQPAQDTTTSGVSEQYNPTTTGISPKAQTMQGQQAIVPVTTPVTTTFTPTNPTNRSKRESKVQIILTFPQGLKKYYSKILKLLHLTQE
                    .. .      :    . :    ... : ..     :                :  :        *** :*: . :****:.*******.::*:**:** *
Prim.cons.          PA3SRPPTPPTTAQ33TENNSH2TPEQ3ES22Q3ATP2PMTSPTQQ2LP2SATPIT2QD2HPSPTNRSKRNL22EIILTLSQGLKKYYGKILKLLHLTLE

                           310       320       330       340       350       360       370       380       390       400
                             |         |         |         |         |         |         |         |         |         |
Q8JSZ3              EDTEGLLEWCKRNLGLDCDDTFFQKRIEEFFITGEGHFNEVLQFRTPGTLSTTESTPAGLPTAEPFKSYFAKGFLSIDSGYYSAKCYSGTSNSGLQLINI
Q70UR1_9VIRU        EDTEGLLEWCKRNLGLDCDDTFFQKRIEEFFITGEGHFNEVLQFRTLGTLSTTESTHAGSPTVEPFKSYFAKGFLSIDSGYFSAKCYSRTSNSGLQLINV
A0A345K523_9VIRU    EDTEGLLEWCKRNLGSNCDDDFFQKRIEEFFITGEGYFNEVLQFKTLSTPSSTEPSHARLPTAEPFKSYFAKGFLSIDSGYFSAKCYPRSSTSGLQLINV
Q0P0I9_9VIRU        EDTEGLLEWCKRNLGSNCDDDFFQKRIEEFFMTGEGYFNEVLQFKTLSTLSPTEPSHARLPTAEPFKSYFAKGFLSIDSGYFSAKCYPRSSASGLQLINV
A0A5K1KE88_9VIRU    EDTEGLLEWCKRNLGLDCDDTFFQKRIEEFFITGEGHFNEVLQFRTPGTLSTKESTHAGSPTAEPFKSYFAKGFLSIDSGYFSAKCYSRTSNSGLQLINV
A0A1V0G0E6_9VIRU    EDTEGLQEWCVRVLGQTCDDSYFHKRLAEFFITGTGFFNEVLLFRSPSTPTTIKPSPTGSPTVEPFRSYYAKGSLTIDSGYFSAKCYPRTSNSGLQLINL
Q8QZF9_9VIRU        EDSEGLLEWCTRVLEQACDDDYFNKRIEEFFITGKGYFNDVLQFRLHDALSNTEPAQTTSPTAVPFKSYYAKGSLTFDSGYFSARCYPRASNSGLQLINV
A0A482LVV7_9VIRU    EDSEGLLEWCTRVLEQACDDDYFNKRIEEFFITGKGYFNDVLQFRLHDAISNTEPAQTTSPTAVPFKSYYAKGSLTFDSGYFSARCYPRASNSGLQLINV
A0A1L3HE22_9VIRU    EDSEGLLEWCTRVLQQVCDDTYFNERIEEFFITGKGYFNDVLQFKLHDTPTTTVSALAASSTAMPFRSYYAKGSLTLDSGYFSAKCYPRASNSGMQLINV
                    **:*** *** * *   *** :*::*: ***:** *.**:** *:  .: :   .: :  .*. **:**:*** *::****:**:**. :* **:****:
Prim.cons.          EDTEGLLEWCKRNLGQDCDD2FFQKRIEEFFITGEGYFNEVLQFRT33TLSTTEP3HAGSPTAEPFKSYFAKGFLSIDSGYFSAKCYPRTSNSGLQLINV

                           410       420       430       440       450       460       470       480       490       500
                             |         |         |         |         |         |         |         |         |         |
Q8JSZ3              TRHSTRIVDTPGPKITNLKTINCINLKASIFKEHREVEINVLLPQVAVNLSNCHVVIKSHVCDYSLDIDGAVRLPHIYHEGVFIPGTYKIVIDKKNKLND
Q70UR1_9VIRU        TRHSTRIADTPGPKITNLKTINCMNLKASVFKEHREVEINVLLPQVAVNLSNCHVAIKSHICDYSLDTDGAIRLPQIHHEGTFIPGTYKIVIDKKSRLND
A0A345K523_9VIRU    TQHPARIAETPGPKTTSLKTINCINLRASVFKEHREVEINVLLPQIAVNLSNCHVVINSHVCDYSLDTDGPVRLPRIYHEGTFMPGTYKIVIDRKNKLND
Q0P0I9_9VIRU        TQHPARIAETPGPKTTSLKTINCINLRASVFKEHREIEINVLLPQIAVNLSNCHVVINSHVCDYSLDTDGPVRLPRIYHEGTFIPGTYKIVIDKKNKLND
A0A5K1KE88_9VIRU    TRHSTRIADTPGPKITNLKTISCINLKASVFKEHREVEINVLLPQVAVNLSNCHVTIKSHVCDYSLDIDGTVRLPHIHHDGTFIPGTYKIVIDRKNKLND
A0A1V0G0E6_9VIRU    TQHSVEIANTPGPKHSSLKTINCINIKASTDKEHNEVEINVLLPQVAINLSNCHVSIRSHVCDYSLDTDGKIRLPQISHEGTFIPGTYKIIIDKKNKLND
Q8QZF9_9VIRU        TQHPVKIANTPGPKLSNLKTINCINLKVSTDKEHSELEVNVLLPQVAVNLSNCHVSIKSHVCDYSLDTDGAIKLPRVAHNGVFIPGTYKIVIDKKNNLND
A0A482LVV7_9VIRU    TQHPVKIANTPGPKLSNLKTINCINLKVSTDKEHSELEVNVLLPQVAVNLSNCHVSIKSHVCDYSLDTDGAIKLPRVAHNGVFIPGTYKIVIDKKNNLND
A0A1L3HE22_9VIRU    TQHPMKIANTPGPKTSNLKTMNCINLKVSTDKEHSELEINVLIPQVAVNLSNCLVSIKSHVCDYSLDTDGAIRLPKVAYNGVFMPGTYKIVIDKKNRLND
                    *:*. .*.:***** :.***:.*:*::.*  *** *:*:***:**:*:***** * *.**:****** ** ::**:: ::*.*:******:**:*..***
Prim.cons.          TQHP2RIANTPGPK2TNLKTINCINLKAS2FKEHREVEINVLLPQVAVNLSNCHVSIKSHVCDYSLDTDGAIRLPRI2HEGTFIPGTYKIVIDKKNKLND

                                                       Gn    1         10        20        30        40        50
                                                             |         |         |         |         |         |
                           510       520       530       540       550       560       570       580       590       600
                             |         |         |         |         |         |         |         |         |         |
Q8JSZ3              RCTLFTDCVIKGREVRKGQSVLRQYKTEIRIGKASTGSRRLLSEEPSDDCISRTQLLRTETAEIHGDNYGGPGDKITICNGSTIVDQRLGSELGCYTINR
Q70UR1_9VIRU        RCTLFTNCVIKGREVRKGQSVLRQYKTEIRIGRASTGSRRLLSEESGDDCISRTQLLRTETAEVHGDNYGGPGDKITICNGSTVVDQRLGSELGCYTINR
A0A345K523_9VIRU    RCTLVTNCVIKGREVRKGQSVLRQYKTEIKIGKAPTGSRKLLSEEPGDDCISRTQLLRTETAEIHDDNYGGPGDKITICNGSTIVDQRLGSELGCYTINR
Q0P0I9_9VIRU        RCTLVTNCVIKGREVRKGQSVLRQYKTEIKIGKASTGFRKLLSEEPGDDCISRTQLLRTETAEIHDDNYGGPGDKITICNGSTIVDQRLGSELGCYTINR
A0A5K1KE88_9VIRU    RCTLFTNCVIKGREVRKGQSVLRQYKTEIRIGKASTGSRRLLSDESGDDCISRTQLLRTETAEIHGDNYGGPGDKITICNGSTIVDQRLGSELGCYTINR
A0A1V0G0E6_9VIRU    RCSLITSCVIKGRELRKGQSTLRQYKTEIKIGQTSSGSRRLLSESSASGCISRTQMIKSETAEIHDDIYGGPGEKLTICNGTTVVDQRLGSELGCYTINR
Q8QZF9_9VIRU        RCTLITNCVIKGREVRKGQSALRQYKTEIRIGQTFTGSRRLLAEGGSSDCISRTQLVKTEVAEIHEDSYGGPGEKITICNGSTIVDQRLGSELGCYTINR
A0A482LVV7_9VIRU    RCTLTTNCVIKGREVRKGQSALRQYKTEIRIGQTVTGSRRLLAEGGSSDCISRTQLVKTEVAEIHEDSYGGPGEKITICNGSTIVDQRLGSELGCYTINR
A0A1L3HE22_9VIRU    RCALITNCVIKGREVRKGQSTLRQYKTEIRIGQTLMGSRRLLAEGVSNDCVSRTQLIKTEVAEIHEDSYGGPGEKITICNGSTIVDQRLGSELGCYTINR
                    **:* *.*******:*****.********:**::  * *:**::  ...*:****::::*.**:* * *****:*:*****:*:****************
Prim.cons.          RCTL2TNCVIKGREVRKGQSVLRQYKTEIRIG2ASTGSRRLLSEE22DDCISRTQLLRTETAEIH3DNYGGPGDKITICNGSTIVDQRLGSELGCYTINR

                    60        70        80        90       100       110       120       130       140       150
                     |         |         |         |         |         |         |         |         |         |
                           610       620       630       640       650       660       670       680       690       700
                             |         |         |         |         |         |         |         |         |         |
Q8JSZ3              VRSFKLCENSATGKNCEIDSVPVKCRQGYCLRITQEGRGHVKLSRGSEVVLDACDTSCEIMIPKGTGDILVDCSGGQQHFLKDNLIDLGCPKIPLLGKMA
Q70UR1_9VIRU        VRSFKLCENSATGKSCEIDSIPVKCKQGYCLKITQEGRGHVKLSRGSEVVLDVCDSSCEVMIPKGTGDILVDCSGGQQHFLKDNLVDLGCPKIPLLGKMA
A0A345K523_9VIRU    VKSFKLCENSATGKTCEIDSTPVKCRQGFCLKITQEGRGHVKLSRGSEVVLDACDSNCEVMIPKGTGDILVDCSGGQQHFLKDNLIDLGCPHIPLLGKMA
Q0P0I9_9VIRU        VKSFKLCENSATGKTCEVDSTPVKCRQGFCLKITQEGRGHVKLSRGSEVVLDACDSSCEVMIPKGTGDILVDCSGGQQHFLKDNLIDLGCPHIPLLGKMA
```

```
A0A5K1KE88_9VIRU   VKSFKLCENSATGKSCEIDSIPVKCRQGYCLKITQEGRGHVKLSRGSEVVLDACDSSCEVMIPKGTGDILVDCSGGQQHFLKDNLIDLGCPKIPLLGKMA
A0A1V0G0E6_9VIRU   IKTYKLCENSVKEKTCEIDSVPVKCRQGLCLKITQEGRGHVKLSRGSEIVLDICDSSCELMIPKGTGDILVDCSGGQQHFLQDNLVDLGCPNIPLLGKVA
Q8QZF9_9VIRU       VRSYKLCENSASGKSCEIDSIPVKCKQGFCLKITQEGRGHVKLSRGTEIVLDVCDSSCELMIPKGTGDILVDCSGGQQHFLQNNLIDLGCPNVPLLGKMA
A0A482LVV7_9VIRU   VKSYKLCENSASGKSCEIDSIPVKCKQGFCLKITQEGRGHVKLSRGTEIVLDVCDSSCELMIPKGTGDILVDCSGGQQHFLQNNLIDLGCPNVPLLGKMA
A0A1L3HE22_9VIRU   VKTYKLCENSATEKGCEIDSVPVKCKQGFCLKVTQEGRGHVKLSRGTEIVLDVCDTSCELMIPKGTGDILVDCSGGQQHFLQDNLIDLGCPNVPLLGKMA
                   ::::******.. * **:** ****:** **::*************:*:*** **:.**:*********************::**:*****::*****:*
Prim.cons.         VKSFKLCENSATGKSCEIDSIPVKCRQGFCLKITQEGRGHVKLSRGSEVVLD2CDSSCE2MIPKGTGDILVDCSGGQQHFLKDNLIDLGCPNIPLLGKMA

                  160       170       180       190       200       210       220       230       240       250
                    |         |         |         |         |         |         |         |         |         |
                          710       720       730       740       750       760       770       780       790       800
                            |         |         |         |         |         |         |         |         |         |
Q8JSZ3             IYICRMSNHPKTTMAFLFWFSFGYVITCILCKAIFYLLIIVGTLGKRLKQYRELKPQTCTICETTPVNAIDAEMHDLNCSYNICPYCASRLTSDGLARHV
Q70UR1_9VIRU       IYICRMSNHPKTTMAFLFWFSFGYVITCILCKAIFFLLIIFGTLGKRFKQYRELKPQTCTICETTPVNAIDAEMHDLNCSYNICPYCASRLTSDGLARHV
A0A345K523_9VIRU   IYICRMSNHPKTTMAFLFWFSFGYVITCIFCKALFYSLIIIGTLGKKIKQYRELKPQTCTICETAPVNAIDAEMHDLNCSYNICPYCASRLTSDGLARHV
Q0P0I9_9VIRU       IYICRMSNHPRTTMAFLFWFSFGYVITCISCKALFYSLIIIGTLGKKIKQYRELKPQTCTICETAPVNAIDAEMHDLNCSYNICPYCASRLTSDGLARHV
A0A5K1KE88_9VIRU   IYICRMSNHPKTTMAFLFWFSFGYVITCIVCKAIFYLLIIVGTLGKRFKQYRELKPQTCTICETTPTNAIDAEMHDLNCSYNICPYCASRLTSDGLARHV
A0A1V0G0E6_9VIRU   IYICRMSNHPKTTMAFLFWFSFGYVITCILCKILFYLLIITGTLGKKLKQYRELKPQTCTICESTPVNAIDAEMHDLNCSYNICPYCASRLTSDGLIRHV
Q8QZF9_9VIRU       IYVCRMSNHPKTTMAFLFWFSFGYVITCIACRVLFYLLIAIGTVGKKLKQYRELKPQTCIVCESVPVNAIDAEMHELNCSYNICPYCASRLSTDGLIRHV
A0A482LVV7_9VIRU   IYVCRMSNHPKTTMAFLFWFSFGYVITCIVCRVLFYLLIAIGTVGKKLKQYRELKPQTCIVCESVPVNAIDAEMHELNCSYNICPYCASRLSTDGLIRHV
A0A1L3HE22_9VIRU   IYICRMSNHPKTTMAFLFWFSFGYVITCVVCKVLFYLLIVIGTLGKKLKQYRELKPQTCIVCESIPVNVIDAEMHELNCNYNICPYCASRLTSDGLVRHV
                   **:*******:*****************: *: :*: **  **:**::*********** :**: *.*.******:***.***********::*** ***
Prim.cons.         IYICRMSNHPKTTMAFLFWFSFGYVITCI2CKALFYLLIIIGTLGKKLKQYRELKPQTCTICETTPVNAIDAEMHDLNCSYNICPYCASRLTSDGLARHV

                  260       270       280       290       300       310        NSm  1        10        20        30
                    |         |         |         |         |         |             |         |         |
                          810       820       830       840       850       860       870       880       890       900
                            |         |         |         |         |         |         |         |         |         |
Q8JSZ3             IQCPKRKEKVEETELYLNLERIPWVVRKLLQVSESTGVALKRSSWLIVLLVLFTVSLSPVQSAPIGQGKTIEAYRAREGYTSICLFVLGSILFIVSCLMK
Q70UR1_9VIRU       TQCPRRKEKVEETELYLNLERIPWVVRKLLQVSESTGTVLKRSSWLIVLLVLFTVSLSPVQSAPIGHGKTIETYRVREEYTSICLFVLGSILFIVSFLMK
A0A345K523_9VIRU   KQCPKRKEKVEETELYLNLERIPWVVRKLLQVSESTGVALKRSSWLIVLLVLLTVSLSPVQSAPVGHGKTIETYQTREGFTSICLFMLGSILFIVSCLVK
Q0P0I9_9VIRU       TQCPKRKEKVEETELYLNLERIPWIVRKLLQVSESTGVALKRSSWLIVLLVLLTVSLSPVQSAPVGHGKTIEIYQTREGFASICLFMLGSILFIVSCLVK
A0A5K1KE88_9VIRU   THCPKRKEKVEETELYLNLERIPWVVRKLLQVSESTGVALKRSSWLIVLLVLFAISLSPVQSAPISHGKTIEIYQVREEYTSICLFVLGSILFIVSCLMK
A0A1V0G0E6_9VIRU   NLCPKRKEKVEETELYLNLERIPWFVRKLLQVSESTGVALKRSSWLLILVILLTVSMSPVQSAPINQQRAVEAYQAREGYTGICLFILGSVLFMTSWLTK
Q8QZF9_9VIRU       TQCPKRREKVEETELFLNLERIPWLVRKLLQVSESTGMALKRSCWMITLLILLLVSISPVQSAPVGKERAIETYQARETYTGICLFVLGSVLFAVSWLTK
A0A482LVV7_9VIRU   TQCPKRREKVEETELFLNLERIPWLVRKLLQVSESTGMALKRSCWMITLLILLLVSMSPVQSAPVGKERAIETYQAREAYTGICLFVLGSVLFAVSWLTK
A0A1L3HE22_9VIRU   TQCPKRKEKVEETELYLNLERIPWLVRRLLQVSESTGMALKRSCWMTTLLILLVVSLSPVQSAPVGQEKAIEVYQVRESYTSMCLFVLGSVLFAVSWLVK
                     **:*:*********:********.**:*********  .****.*:  *::*: :*:*******:.: :::* *:.** ::.:***:***:** .* * *
Prim.cons.         TQCPKRKEKVEETELYLNLERIPWVVRKLLQVSESTGVALKRSSWLIVLLVLLTVSLSPVQSAPVGHGKTIETYQAREGYTSICLFVLGSILFIVS2L3K

                          40        50        60        70        80        90       100       110       120       130
                            |         |         |         |         |         |         |         |         |         |
                          910       920       930       940       950       960       970       980       990      1000
                            |         |         |         |         |         |         |         |         |         |
Q8JSZ3             GLVDSVGNSFFPGLSICKTCSISSINGFEIESHKCYCSLFCCPYCRHCSTDKEIHKLHLSICKKRKKGSNVMLAVCKLMCFRATMEVSNRALFIRSIINT
Q70UR1_9VIRU       GLVDGVGNIFFPGLSVCKTCSIGSINGFEIESHKCYCSLFCCPYCRHCSADGEIHQLHLSICKKRKTGSNVMLAVCKRMCFRATMEVSNKALFIRSIINT
A0A345K523_9VIRU   GLVDSVSDSFFPGLSVCKTCSIGSINGFEIESHKCYCSLFCCPYCRHCSADREIHQLHLSICKKRKTGSNVMLAVCKRMCFRATIEASRRALLIRSIINT
Q0P0I9_9VIRU       GLVDSVSDSFFPGLSVCKTCSIGSVNGFEIESHKCYCSLFCCPYCRHCSADREIHQLHLSICKKRKTGSNVMLAVCKRMCFRATIEASRRALLIRSIINT
A0A5K1KE88_9VIRU   GLVNSVGNIFFPGLSICKTCSIGSINGFEIESHKCYCSLLCCPYCRHCSADREIHQLHLSICKKRKTGSNVMLAVCKRMCFKATMEVSNKALFIRSIINT
A0A1V0G0E6_9VIRU   GLIDSIGDSFFPGLFVCKTCSIGSINGFEIESHRCYCSLLCCPYCRTCSADKDSHQLHLSICKKKKAGSNVMLAVCKRMCFKATIEASNRVLLIRSIINT
Q8QZF9_9VIRU       ALIDGIGNSFFPGLSVCKTCSIGSINGFEIESHKCYCSLFCCPYCRACSSDKNTHRMHLNVCKKRKMGSNVMLAVCKRMCFRATIEASNKALLIRSIINS
A0A482LVV7_9VIRU   ALIDGIGNSFFPGLSVCKTCSIGSINGFEIESHKCYCSLFCCPYCRACSSDKNTHRMHLNVCKKRKMGSNVMLAVCKRMCFRATIEASNKALLIRSIINS
A0A1L3HE22_9VIRU   GLIDGIGNSFFPGLSVCKTCSIGSINGFEIESHKCYCSLFCCPYCRACSSDKITHRMHLNICKRRKVGSNVMLAVCKRMCFKATVEASNSALLIRSIINS
                   .*::.:.: ***** :******.*:*********:*****:****** **:*   *::**.:**::* ********** ***:**:*.*.  .*:******:
Prim.cons.         GLVDSVGNSFFPGLSVCKTCSIGSINGFEIESHKCYCSLFCCPYCRHCSADKEIHQLHLSICKKRKTGSNVMLAVCKRMCFRATIEASN2ALLIRSIINT

                         140       150
                            |         |
                         1010      1020      1030      1040      1050      1060      1070      1080      1090      1100
                            |         |         |         |         |         |         |         |         |         |
Q8JSZ3             TFVLCILILAVCVV-STSAVEMENLPAGTWEREEDLTNFCHQECQVTETECLCPYEALVLRKPLFLDSTAKGMKNLLNSTSLETSLSIEAPWGAINVQST
Q70UR1_9VIRU       TFVMCILKGEFAAVKSIHPIEMESLPAGTWEREEDLTNFCHQECQVTETECLCPYEALVLRRPLFLDSIVKGMKNLLNSTSLETSLSIEAPWGAINVQST
A0A345K523_9VIRU   TFVICILTLTICVV-STSAVEMENLPAGTWEREEDLTNFCHQECQVTETECLCPYEALVLRKPLFLDSIVKGMKNLLNSTSLETSLSIEAPWGAINVQST
Q0P0I9_9VIRU       TFVICILTLTICVV-STSAVEMENLPAGTWEREEDLTNFCHQECQVTETECLCPYEALVLRKPLFLDSIVKGMKNLLNSTSLETSLSIEAPWGAINVQST
A0A5K1KE88_9VIRU   TFVVCILILAVCVV-STSAVEMENLPAGTWEREEDLTNFCHQECQVTETECLCPYEALVLRRPLFLDSIVKGMKNLLNSTSLETSLSIEAPWGAINVQST
A0A1V0G0E6_9VIRU   TFIICMLILVVCVI-STSAVEMEDLPAGTWEREEDLTNFCHQECQVTETECLCPYEAFVLRKPLFLSSIVKGKKNLLNSASLETSLSIEAPWGAINAQST
Q8QZF9_9VIRU       TFVICILILTICVV-STSAVDMENLPAGIWEKEEDLTNFCHQECQVTETECLCPYEAMVLRKPLFLDNIAKGMKSLLNSTSLETSLSIEAPWGAINVQST
A0A482LVV7_9VIRU   TFVICILILTICVV-STSAVDMENLPAGIWEKEEDLTNFCHQECQVTETECLCPYEAMVLRKPLFLDSIAKGMKSLLNSTSLETSLSIEAPWGAINVQST
A0A1L3HE22_9VIRU   TFVICVLILVICVV-STSAVDMENLPAGIWEREEDLTNFCHQECQVTETECLCPYEALILRKPLYLDSIAKGMKSLLNSTNLETSLSIEAPWGAINVQST
                   **::*:*   ...: *  .::**.**** **:***************************::**:**:*.. .** *.****:.**************.***
Prim.cons.         TFVICILILTICVVKSTSAVEMENLPAGTWEREEDLTNFCHQECQVTETECLCPYEALVLRKPLFLDSIVKGMKNLLNSTSLETSLSIEAPWGAINVQST

                         1110      1120      1130      1140      1150      1160      1170      1180      1190      1200
                            |         |         |         |         |         |         |         |         |         |
Q8JSZ3             YKPTVSTANIALSWSSVEHRGNKILVSGRSESIMKLEERTGISWDLGVEDASESKLLTVSVMDLSQMYSPVFEYLSGDRQVGEWPKATCTGDCPERCGCT
Q70UR1_9VIRU       YKPTVSTANIALSWSSVEHRGNKVLVSGRSESIMKLEERTGISWDLGVEDASESKLLTVSVMDLSQMYSPVFEYLSGDRQVEEWPKATCTGDCPERCGCT
A0A345K523_9VIRU   FKPTVSTANIALSWSSVEHRGNKILVTGRSESIMKLEERTGVSWDLGVEDASESKLLTVSIMDLSQMYSPVFEYLSGDRQVEEWPKATCTGDCPQRCGCT
Q0P0I9_9VIRU       FKPAVSTANIALSWSSVEHRGNKILVTGRSESIMKLEERTGVSWDLGVEDASESKLLTVSIMDLSQMYSPVFEYLSGDRQVEEWPKATCTGDCPERCGCT
A0A5K1KE88_9VIRU   YKPTVSAANIALSWSSVEHRGNKILVSGRSESIMKLEERTGISWDLGVEDASESKLLTVSVMDLSQMYSPVFEYLSGDRQVEEWPKATCTGDCPERCGCT
A0A1V0G0E6_9VIRU   FKPTISTANIALSWSSVEHRGNKVLVSGRSESIMKLEERTGISWNLGVEDASESRMLTVSVMDLSQMYLPVFEYLSGDRQVEEWPKATCTGDCPERCGCT
Q8QZF9_9VIRU       FKPTVSAANIALSWSSVEHRGNKILVSGRSESIMKLEERTGVSWNLGVEDASESKTLTVSVMDLSQMYSPVFEYLSGDRQVEEWPKATCTGDCPERCGCT
A0A482LVV7_9VIRU   FKPTVSAANIALSWSSVEHRGNKILVSGRSESIMKLEERTGVSWNLGVEDASESKTLTVSVMDLSQMYSPVFEYLSGDRQVEEWPKATCTGDCPERCGCT
A0A1L3HE22_9VIRU   FKPTVSAANIALSWSSVEHRGNKILVSGRSESIMKLEERTGISWSLGVEDASESKTLTVSVMDLSQMYSPVFEYLSGDRQVEEWPKATCTGDCPERCGCT
                   :**::*:****************:**:**************:**.*********: ****:******* ************ ************:*****
Prim.cons.         FKPTVSTANIALSWSSVEHRGNKILVSGRSESIMKLEERTGISWDLGVEDASESKLLTVSVMDLSQMYSPVFEYLSGDRQVEEWPKATCTGDCPERCGCT

                         1210      1220      1230      1240      1250      1260      1270      1280      1290      1300
                            |         |         |         |         |         |         |         |         |         |
Q8JSZ3             SSTCLHKEWPHSRNWRCNPTWCWGVGTGCTCCGLDVKDLFTDYMFVKWKVEYIKTEAIVCVELTSQERQCSLIEAGTRFNLGPVTITLSEPRNIQQKLPP
Q70UR1_9VIRU       SSTCLHKEWPHSRNWRCNPTWCWGVGTGCTCCGLDVKDLFTDYMFVKWKVEYIKTEAIVCVELTSQERQCSLIEAGTRFNLGSVTITLSEPRNIQQKLPP
A0A345K523_9VIRU   SSTCLHKEWPHSRNWRCNPTWCWGVGTGCTCCGVDVKDLFTDHMFVKWKVEYIKTEAIVCVELTSQERQCSLIEAGTRFNLGPVTITLSEPRNIQQKLPP
Q0P0I9_9VIRU       SSTCLHKEWPHSRNWRCNPTWCWGVGTGCTCCGVDVKDLFTDHMFVKWKVEYIKTEAIVCVELTSQERQCSLIEAGTRFNLGPVTITLSEPRNIQQKLPP
A0A5K1KE88_9VIRU   SSTCLHKEWPHSRNWRCNPTWCWGVGTGCTCCGLDVKDLFTDYMFVKWKVEYIKTEAIVCVELISQERQCSLIEAGTRFNLGPVTITLSEPRNIQQKLPP
A0A1V0G0E6_9VIRU   SSTCLHKEWPHSRNWRCNPTWCWGVGTGCTCCGLDVKDLFTDYMFVKWKVEYIKTEAIVCVELTSQERQCSLIEAGTRFNLGPVTITLSEPRNIQQKLPS
Q8QZF9_9VIRU       SSTCLHKEWPHSRNWRCNPTWCWGVGTGCTCCGLDVKDFFTDYMLVKWKVEYIKTEAIVCVELTSQERQCSLIEAGTRFNLGPVTITLSEPRNVQQRLPP
A0A482LVV7_9VIRU   SSTCLHKEWPHSRNWRCNPTWCWGVGTGCTCCGLDVKDFFTDYMLVKWKVEYIKTEAIVCVELTSQERQCSLIEAGTRFNLGPVTITLSEPRNVQQRLPP
A0A1L3HE22_9VIRU   SSTCLHKEWPHSRNWRCNPTWCWGVGTGCTCCGLDVRDLFTDYMMVKWKVEYVKTEAIMCVELTSQERQCSLIEAGTRFNLGPVTITLSEPRNVQQRLPS
                   ********************************:**:*:***:*:*******:*****:**** *****************.**********:**:**.
Prim.cons.         SSTCLHKEWPHSRNWRCNPTWCWGVGTGCTCCGLDVKDLFTDYMFVKWKVEYIKTEAIVCVELTSQERQCSLIEAGTRFNLGPVTITLSEPRNIQQKLPP

                         1310      1320      1330      1340      1350      1360      1370      1380      1390      1400
```

```
                          |         |         |         |         |         |         |         |         |         |
Q8JSZ3             EIITLHPRIEEGFFDLMHVQKVLSASTVCKLQSCTHGVPGDLQVYHIGNLLKGDKVNGHLIHKIEPHFNTSWMSWDGCDLDYYCNMGDWPSCTYTGVTQH
Q70UR1_9VIRU       EIITLHPKIEEGFFDLMHVQKVLSASTVCKLQSCTHGVPGDLQVYHIGNLLKGDRVNGHLIHKIEQHFNTSWMSWDGCDLDYYCNMGDWPSCTYTGVTQH
A0A345K523_9VIRU   EIITLHPKIEEGFFDLMHVQKVLSASTVCKLQSCTHGIPGDLQVYHIGNLLKGDRVNGHLIHKIEPYFNTSWMSWDGCDLDYYCNMGDWPSCTYTGVTQH
Q0P0I9_9VIRU       EIITLHPKIEEGFFDLMHVQKVLSASTVCKLQSCTHGIPGDLQVYHIGNLLKGDRVNGHLIHKIESHFNTSWMSWDGCDLDYYCNMGDWPSCTYTGVTQH
A0A5K1KE88_9VIRU   EIITLHPRIEEGFFDLMHVQKVLSASTVCKLQSCTHGVPGDLQVYHIGNLLKGDKVNGHLIHKIEPHFNTSWMSWDGCDLDYYCNMGDWPSCTYTGVTQH
A0A1V0G0E6_9VIRU   EIVTLHPKIEEGFFDLMHVQKVLSASTVCKLQSCTHGVAGDLQVYHVGNLLKGDKINGHSIHKIEPHLNTSWMSWDGCDLDYYCNMGDWPSCTYTGVTQH
Q8QZF9_9VIRU       EIVTLHPKIEEGFFDLMHVQKILSASTVCKLQSCTHGVPGDFQVYHVGNLLRGDKVNGHSIHKIEPHFNTSWMSWDGCDLDYFCNMGDWPSCTYTGVTQH
A0A482LVV7_9VIRU   EIVTLHPKIEEGFFDLMHVQKILSASTVCKLQSCTHGVPGDFQVYHVGNLLRGDKVNGHSIHKIEPHFNTSWMSWDGCDLDYFCNMGDWPSCTYTGVTQH
A0A1L3HE22_9VIRU   EIVTLHPKIEEGFFDLMHVQKVLFCKHSLQATSCTHGVPGDLQVYHVGNLLKGDKVNGHSIHKIESHFNTSWMSWDGCDLDYFCNMGDWPSCTYTGVTQH
                   **:****:*************:*  ..   :  *****:.**:****:****:**::*** *****  ::**************:*****************

Prim.cons.         EIITLHPKIEEGFFDLMHVQKVLSASTVCKLQSCTHGVPGDLQVYHIGNLLKGDKVNGHLIHKIEPHFNTSWMSWDGCDLDYYCNMGDWPSCTYTGVTQH

                         1410      1420      1430      1440      1450      1460      1470      1480      1490      1500
                          |         |         |         |         |         |         |         |         |         |
Q8JSZ3             NHASFVNLLNIETDYTKNFHFHSKRVTAHGDTPQLDLKARPTYGAGEITVLVEVADMELHTKKIEISGLKFASLACTGCYACSSGISCKVRIHVDEPDEL
Q70UR1_9VIRU       NHASFVNLLNIETDYTKTFHFHSKRVTAHGDTPQLDLKARPTYGAGEITVLVEVADMELHTKKIEISGLKFAGLTCTGCYACSSGISCKVRIHVDEPDEL
A0A345K523_9VIRU   NHAAFVNLLNIETDYTKTFHFHSKRVTAHGDTPQLDLKARPTYGAGEITVLVEVADMELHTKKVEISGLKFASLACTGCYACSSGISCKVRIHVDEPDEL
Q0P0I9_9VIRU       NHAAFVNLLNIETDYTKTFHFHSKRVTAHGDTPQLDLKARPTYGAGEITVLVEVADMELHTKKVEISGLKFASLACTGCYACSSGISCKVRIHVDEPDEL
A0A5K1KE88_9VIRU   NHASFVNLLNIETDYTKSFHFHSKRVTAHGDTPQLDLKARPNYGAGEITVLVEVADMELHTKKIEISGLKFASLTCTGCYACSSGISCKVRIHVDEPDEL
A0A1V0G0E6_9VIRU   NHAAFVNLLNIETDYTKSFHFHSKRVTAHGDTPQLDLKARPMYGAGEITVLVEVADMELHSKNIEISGLKFASLTCSGCYACSSGISCKVRIHVDEPDEL
Q8QZF9_9VIRU       NRAAFINMLNIETDYTKTFHFHSKRVTAHGDTPQLDLKARPAYGAGEVTVLVEVADLELHTKKLEVSGLKIASLTCSGCYACSSGISCKVRIHVNEPDEF
A0A482LVV7_9VIRU   NRAAFINMLNIETDYTKTFHFHSKRVTAHGDTPQLDLKARPAYGAGEVTVLVEVADLELHTKKLEVSGLKIASLTCSGCYACSSGISCKVRIHVNEPDEF
A0A1L3HE22_9VIRU   NRAAFVNMLNIETDYTKTFHFHSKRVTAHGDTPQLDLKARPTYGAGEVTVLVEVADLELHTKKIEVSGLKFASLTCSGCYACSSGISCKVRIHVNEPDEF
                   *:*:*:*:*********.*********************** *****:********:***:*::*:****:*.*:*:****************:****:

Prim.cons.         NHAAFVNLLNIETDYTKTFHFHSKRVTAHGDTPQLDLKARPTYGAGEITVLVEVADMELHTKKIEISGLKFASLTCTGCYACSSGISCKVRIHVDEPDEL

                         1510      1520      1530      1540      1550      1560      1570      1580      1590      1600
                          |         |         |         |         |         |         |         |         |         |
Q8JSZ3             TVHVKSDDPDVVAASSSLMARKLEFGTDSTFKAFSAMPKTSLCFYIVEREHCKSCSEEDTKKCVNTKLEQPQSILIEHKGTIIGKQNSTCTAKASCWLES
Q70UR1_9VIRU       TVHVKSDDPDVVAASSSLMARKLEFGTDSTFKAFSAMPKTSLCFYIVEREYCKSCNKEDTQKCVNTKLEQPQSILIEHKGTIIGKQNNTCTAKASCWLES
A0A345K523_9VIRU   TVHVKSSDPDVVAAGTSLMARKLEFGTDSTFKAFSAMPKTSLCFYIVEKEYCKSCSEDDTKKCVDTKLEQPQSILIEHRGTIIGKQNDTCTAKASCWLES
Q0P0I9_9VIRU       TVHVKSSDPDVVAASTSLMARKLEFGTDSTFKAFSAMPKTSLCFYIVEREYCKSCSEDDTQKCVDTRLEQPQSILIEHKGTIIGKQNDTCTAKASCWLES
A0A5K1KE88_9VIRU   TVHVKSDDPDVVAASSSLMARKLEFGTDSTFKAFSAMPKTSLCFYIVEREYCKSCSKEDTQKCVNTKLEQPQSILIEHKGTIIGKQNNTCTSKTSCWLES
A0A1V0G0E6_9VIRU   AVHVKSDDLDVVAASSSLMARKIEFGTDSTFKAFSSMPKESLCFYIVERDFCKSCAAEDTQKCVKAKLEQPQSILIEHKGTIIGKQNDTCSTKTSCWLES
Q8QZF9_9VIRU       TVHVRSSDPDVVAAGSSLMARKIEFGADSTFKAFSSMPKDSLCFYIVEKDYCSSCTDDDTQKCVNTKLDHPQSILIEHKGTIIGKQNDTCSTKTSCWLES
A0A482LVV7_9VIRU   TVHVRSSDPDVVAAGSSLMARKIEFGADSTFKAFSSMPKDSLCFYIVEKDYCSSCTDDDTQKCVNTKLDHPQSILIEHKGTIIGKQNDTCSTKTSCWLES
A0A1L3HE22_9VIRU   TVHVKSNDPDVVAAGSSLMARKIELGADSTFKAFSSMPKNSLCFYIVERDYCSSCTDDDTQKCVNTKLEHPQSILIEHKGTIIGKQNDTCPTKTSCWLES
                   :***:*.* *****.:******:*:*:********:*** ********::.*.**  :**:***.::*::********:********.**.:*:******

Prim.cons.         TVHVKS2DPDVVAASSSLMARKLEFGTDSTFKAFSAMPKTSLCFYIVEREYCKSCS2DDTQKCVNTKLEQPQSILIEHKGTIIGKQNDTCT2KTSCWLES

                         1610      1620      1630      1640      1650      1660      1670      1680      1690      1700
                          |         |         |         |         |         |         |         |         |         |
Q8JSZ3             VKSFFYGLKNMLSGIFGNVFMGIFLFLAPFILLILFFMFGWRILFCFKCCRRTRGLFKYRHLKDDEETGYRRIIEKLNNKK-GKNKLLDGERLADRRIAE
Q70UR1_9VIRU       VKSFFYGLKNMLGGIFGNVFIGIFTFLTPFILLILFFMFGWRILFCFKCCRRTRGLFKYRHLKDDEETGYRKIIERLNNKK-GKNRLLDGERLADRKIAE
A0A345K523_9VIRU   VKSFFYGLKNMLGSVFGNFFIGILLFLAPFVLLVLFFMFGWKILFCFKCCRRTRGLFKYRHLKDDEETGYRRIIERLNNKK-GKNRLLDGERLADRKIAE
Q0P0I9_9VIRU       VKSFFYGLKNMLGSVFGNLFIGILLFLAPFVLLVLFFMFGWKILFCFKCCRRTRGLFKYRHLKDDEETGYRRIIERLNSKK-GKNRLLDGDRLADRKIAE
A0A5K1KE88_9VIRU   VKSFFYGLKNMLSSIFGNVFVGIFLFLAPFILLILFFMFGWRILFCFKCCRRTRGLFRYKHLEDDEETGYRRIIERLNNKK-GKNKLLDGERLADRRIAE
A0A1V0G0E6_9VIRU   FKSFFYGLKNMLSGVFGSAIMGILLFLAPFILLILFFFFGWRILFCLRCCRRTRGLFKYKHIKDDEEAGYKKIIEKLNNKKGGTSKLLDGERLADRKIAE
Q8QZF9_9VIRU       VKGFFYGLKNMLSGIFGNVFLGIFLFLAPFVLLIFFFFFGWRLLFCLKCCRRTRGLLKYKHLKDEEEAGYKKIIERLNGKK-GKSQLFDGERLADRKIAE
A0A482LVV7_9VIRU   VKGFFYGLKNMLSGIFGNVFLGILLFLAPFVLLIFFFFFGWRLLFCLKCCRRTRGLLKYKHLKDEEEAGYKKIIERLNGKK-GKSQLFDGERLADRKIAE
A0A1L3HE22_9VIRU   VKSFFYGLKNMLGGIFGNVFLGIFLFLAPFALLILFFFFGWRLLFCLKCCRRTRGLLKYKQLKDEEEAGYKKIIERLNDKK-GKSRLFDGERLADRKIAE
                   .*.*********..:**. ::**: **:** **::**:***::***::********::*::::*:**:**::***:**.** *..:*:**:*****:***

Prim.cons.         VKSFFYGLKNMLSGIFGNVF2GIFLFLAPF2LLILFFMFGWRILFCFKCCRRTRGLFKYKHLKDDEETGYRKIIERLNNKKGGKNRLLDGERLADRKIAE


Q8JSZ3             LFSTKTHIG
Q70UR1_9VIRU       LFSTKTHIG
A0A345K523_9VIRU   LFSTKTHIG
Q0P0I9_9VIRU       LFSTKTHIG
A0A5K1KE88_9VIRU   LFSTKTHIG
A0A1V0G0E6_9VIRU   LFSTKAHIG
Q8QZF9_9VIRU       LFSTKIHIG
A0A482LVV7_9VIRU   LFSTKIHIG
A0A1L3HE22_9VIRU   LFSTKTHIG
                   ***** ***

Prim.cons.         LFSTKTHIG
```

**Table S3: Sequence alignments and numbering of CCHFV and DUGV M polyproteins**. Aligned NSm sequences shown in Figure 3b are highlighted in yellow. Red type/ asterisk (*), green type/colon (:), and blue type/dot (.) indicate identical amino acid residues, conserved substitution, and semi-conserved substitutions respectively.

```
                       10        20        30        40        50        60        70        80        90       100
                        |         |         |         |         |         |         |         |         |         |
Q02004_DUGV    MSKRVLIIAVVVYL-------------------------------------------------------------VF-----------------------
Q8JSZ3_CCHFV   MHISLMYAILCLQLCGLGETHGSHNETRHNKTDTMTTPGDNPSSEPPVSTALSITLDPSTVTPTTPASGLEGSGEVYTSPPITTGSLPLSETTPELPVTT
               *   ::   : : *                                                             *:
Prim.cons.     M222222222222LCGLGETHGSHNETRHNKTDTMTTPGDNPSSEPPVSTALSITLDPSTVTPTTPASGLEGSGEV2TSPPITTGSLPLSETTPELPVTT

                      110       120       130       140       150       160       170       180       190       200
                        |         |         |         |         |         |         |         |         |         |
Q02004_DUGV    ------------TTQNQITGNHTTINSSSPSTTEASSTP------------TVSRTPQTTTTSTAVSTTITATTTP------------TASWTTQSQYFN
Q8JSZ3_CCHFV   GTDTLSAGDVDPSTQTAGGTSAPTVRTSLPNSPSTPSTPQDTHHPVRNLLSVTSPGPDETSTPSGTGKESSATSSPHPVSNRPPTPPATAQGPTENDSHN
                           :**.    . .*:.:* *.:..:.***            ..*  *: *:*.:....  :**::*            **. .*:.: .*
Prim.cons.     GTDTLSAGDVDP2TQ22222222T222S2P222222STPQDTHHPVRNLLS22S22P22T2T222222222AT22PHPVSNRPPTPPATA222T22222N

                      210       220       230       240       250       260       270       280       290       300
                        |         |         |         |         |         |         |         |         |         |
Q02004_DUGV    KTTQH-------------------HWREETMIS---RNPTVLDRQSRASSVRELLNT---------KFLMLLGFIPKGEVNHLENACNRE-GKNCTELIL
Q8JSZ3_CCHFV   ATEHPESLTQSATPGLMTSPTQIVHPQSATPITVQDTHPSPTNRSKRNLKMEIILTLSQGLKKYYGKILRLLQLTLEEDTEGLLEWCKRNLGLDCDDTFF
                * :                    * :. * *:    :*:  :*..*  .:. :*.          *:* ** :  : :.: * : *:*: * :* : ::
Prim.cons.     2T222ESLTQSATPGLMTSPTQIVH2222T2I2VQD22P2222R22R2222222L22SQGLKKYYGK2L2LL2222222222L222C2R2LG22C22222

                      310       320       330       340       350       360       370       380       390       400
                        |         |         |         |         |         |         |         |         |         |
Q02004_DUGV    KERIARFFSETEKESCYNTYLEKHLRSVSPEVSLTPYRVLGLREDILLKE-IDRRIIRFETDSQRVTCLSAS----LLKPDVFIREQRIDAKPSNGPKIV
Q8JSZ3_CCHFV   QKRIEEFF--ITGEGHFNEVLQFRTPGTLSTTESTP---AGLPTAEPFKSYFAKGFLSIDSGYYSAKCYSGTSNSGLQLINITRHSTRIVDTP--GPKIT
               ::** .**     *. :*  *: :  .. . .. **    **     :*. : : :: :::.   ..* *.:    *   ::  :. **  .*  ****.
Prim.cons.     22RI22FFSE222E222N22L2222222222222TPYRV2GL222222K2Y2222222222222222C2S22SNSGL2222222222RI222PSNGPKI2

                      410       420       430       440       450       460       470       480       490       500
                        |         |         |         |         |         |         |         |         |         |
Q02004_DUGV    PVDSVACMNLEANVDVRSNKLVIQSLMTTVKISLKNCKVVVNSRQCIHQQTGSGVIKVPKFEKQQGGTWSSYIAGVYTATIDLLDENNQNCKLFTECIVK
Q8JSZ3_CCHFV   NLKTINCINLKASIFKEHREVEINVLLPQVAVNLSNCHVVIKSHVCDYSLDIDGAVRLPHIYHE-----GVFIPGTYKIVIDKKNKLNDRCTLFTDCVIK
                :.:: *:**:*.:  . .:: *: *:. * :.*.**:**::*: * :.   .*.:::*:: ::     . :*.*.*. .**  :: *:.*.***:*::*
Prim.cons.     222222C2NL2A2222222222I22L222V222L2NC2VV22S22C2222222G2222P22222QGGTW222I2G2Y222ID22222N22C2LFT2C22K

                      510       520       530       540       550       560       570       580       590       600
                        |         |         |         |         |         |         |         |         |         |
Q02004_DUGV    GRELVKGQSELKSFNIEVLLPRVMKTRRKLLAVTDGSTECNSGTQLIEGKSIEVHKQDIGGPGKKLTICNGTSVLDVPLDEGHGCYTINVITSKRACRPK
Q8JSZ3_CCHFV   GREVRKGQSVLRQYKTEIRIGKASTGSRRLLS-EEPSDDCISRTQLLRTETAEIHGDNYGGPGDKITICNGSTIVDQRLGSELGCYTINRVRSFKLCENS
               ***: **** *:.:: *: : :. .  *:**:  : * :* * ***:. :: *:* :: ****.*:*****::::*  *..  ****** : * : *. .
Prim.cons.     GRE22KGQS2L22222E2222222222R2LL2V222S22C2S2TQL222222E2H2222GGPG2K2TICNG2222D22L2222GCYTIN222S222C222

                      610       620       630       640       650       660       670       680       690       700
                        |         |         |         |         |         |         |         |         |         |
Q02004_DUGV    NSKLQCSIDKELKPCDSGKCLSISQKGAGHIKVSRGKTILITECKEHCQIPVPTGKGDIMVDCSGGRQHYLEVNIVDIHCPNTKFLGGIMLYFCRMSSRP
Q8JSZ3_CCHFV   ATGKNCEIDSVPVKCRQGYCLRITQEGRGHVKLSRGSEVVLDACDTSCEIMIPKGTGDILVDCSGGQQHFLKDNLIDLGCPKIPLLGKMAIYICRMSNHP
                :  :*.**.    * .* ** *:*:* **:*:***. :::  *.  *:* :*.*.***:******:**:*: *::*: **:  :** : :*:****.:*
Prim.cons.     22222C2ID22222C22G2CL2I2Q2G2GH2K2SRG2222222C222C2I22P2G2GDI2VDCSGG2QH2L22N22D22CP2222LG2222Y2CRMS22P

                      710       720       730       740       750       760       770       780       790       800
                        |         |         |         |         |         |         |         |         |         |
Q02004_DUGV    TVALLLGIWIGCGYILTCIFSFLLYHLILFFANCIKQCRKKGERLGEICVKCEQQTVNLMDQELHDLNCNFNLCPYCCNRMSDEGMSRHVGKCPKRLERL
Q8JSZ3_CCHFV   KTTMAFLFWFSFGYVITCILCKAIFYLLIIVGTLGKRLKQYRELKPQTCTICETTPVNAIDAEMHDLNCSYNICPYCASRLTSDGLARHVIQCPKRKEKV
               ..:: : :*:. **::***:.  :::*:::...  *: ::  *   : *. **  .** :* *:*****.:*:****..*::.:*::*** :**** *::
Prim.cons.     22222222W222GY22TCI2222222L22222222K222222E22222C22CE222VN22D2E2HDLNC22N2CPYC22R2222G22RHV22CPKR2E22

                      810       820       830       840       850       860       870       880       890       900
                        |         |         |         |         |         |         |         |         |         |
Q02004_DUGV    NEIELYLTTSEC-LCLSVCYQLLISVGIFLKRTTWLVVLLVLLGLAISPVQGAP-----TEVSNVKQDGDYSICYFIFGC-LVTAALLLKVKRTNSNGIV
Q8JSZ3_CCHFV   EETELYLNLERIPWVVRKLLQVSESTGVALKRSSWLIVLLVLFTVSLSPVQSAPIGQGKTIEAYRAREGYTSICLFVLGSILFIVSCLMK----------
               :* ****. ..    :    *:  *.*: ***::**:*****: :::****.**     *  :   ::*  *** *::*. *. .: *:*
Prim.cons.     2E2ELYL22222P2222222Q222S2G22LKR22WL2VLLVL22222SPVQ2APIGQGKT22222222G22SIC2F22G2IL22222L2KVKRTNSNGIV

                      910       920       930       940       950       960       970       980       990      1000
                        |         |         |         |         |         |         |         |         |         |
Q02004_DUGV    VVVDS--------FGRCPYCNEFTDSLFEEVLHDTLCSLCVCPFCEKQALDLVTLEEHVKECYKVATRKDIFKILGRKFTNALVRREKLFTTGLQLFINK
Q8JSZ3_CCHFV   GLVDSVGNSFFPGLSICKTCSISSINGFEIESHKCYCSLFCCPYCRHCSTDKEIHKLHLSICKKRKKGSNVMLAVCK----LMCFRATMEVSNRALFIR-
                :***           *  *.  : . **   *.  ***  **:*.: : *    : *:. * *  . .:::  : :     :  * .: .:.  ***.
Prim.cons.     22VDS2G2SFFPGLSIC22C2222222FE222H222CSL22CP2C22222D222222H222C2K2222222222222KFTN2222R222222222LFI2K

                     1010      1020      1030      1040      1050      1060      1070      1080      1090      1100
                        |         |         |         |         |         |         |         |         |         |
Q02004_DUGV    TNVVVFALIMCFLLL-LTGHNASAFDSGDLPDGVWEESSQLVKSCTQFCYIEEDVCYCPAEDGVGRKLLFFNGLQNSVKRLSDSHKLLTSVSIDAPWGRI
Q8JSZ3_CCHFV   -SIINTTFVLCILILAVCVVSTSAVEMENLPAGTWEREEDLTNFCHQECQVTETECLCPYEALVLRKPLFLDSTAKGMKNLLNSTSLETSLSIEAPWGAI
                .::  ::::*:*:* :   .:**.:  :** *.**...:*.: * * * : *  * ** *  * ** **::.  :.:*.* :* .* **:**:**** *
Prim.cons.     T222222222C2L2LA222222SA22222LP2G2WE2222L222C2Q2C222E22C2CP2E22V2RK2LF22222222K2L22S22L2TS2SI2APWG2I

                     1110      1120      1130      1140      1150      1160      1170      1180      1190      1200
                        |         |         |         |         |         |         |         |         |         |
Q02004_DUGV    NVESTWKPTLAASNIAMSWSSTDIKGEKVILSGRSTSIIKLKEKTGVMWKLVGSGLASEKKKPFRFPIMDFAQVYNSVFQYITGDRLLSEWPKAVCTGDC
Q8JSZ3_CCHFV   NVQSTYKPTVSTANIALSWSSVEHRGNKILVSGRSESIMKLEERTGISWDL-GVEDASESKL-LTVSVMDLSQMYSPVFEYLSGDRQVGEWPKATCTGDC
               **:**:***::::***:****.: :*:*:::**** **:**:*:**: *.* *   ***.*  : ..:**::*:*..**:*::*** :.*****.*****
Prim.cons.     NV2ST2KPT2222NIA2SWSS2222G2K222SGRS2SI2KL2E2TG22W2LVG222ASE2K2P22222MD22Q2Y22VF2Y22GDR222EWPKA2CTGDC

                     1210      1220      1230      1240      1250      1260      1270      1280      1290      1300
                        |         |         |         |         |         |         |         |         |         |
Q02004_DUGV    PHRCGCQTSTCMAKE--------CHTQECVSTHMVLGIGTGCTCCGMDVERPFNKYLGVKWSTEYLRTEVLVCVEVTEEERHCEIVEAGTRFNIGPITIT
Q8JSZ3_CCHFV   PERCGCTSSTCLHKEWPHSRNWRCNPTWC------WGVGTGCTCCGLDVKDLFTDYMFVKWKVEYIKTEAIVCVELTSQERQCSLIEAGTRFNLGPVTIT
               *.**** :***: **        *:.  *       *:********:**:  *..*: ***..**::**.:****:*.:**:*.::*******:**:***
Prim.cons.     P2RCGC22STC22KEWPHSRNWRC2222CVSTHMV2G2GTGCTCCG2DV222F22Y22VKW22EY22TE22VCVE2T22ER2C222EAGTRFN2GP2TIT

                     1310      1320      1330      1340      1350      1360      1370      1380      1390      1400
                        |         |         |         |         |         |         |         |         |         |
```

```
Q02004_DUGV    ISDPQNIGSKLPESLMTVQEIDDSNFVDIMHVGNVISADNSCRLQSCTHGSAVTTRFTALTALIKDDHSSGLNLAVLDPKVNSSWLSWEGCDMDYYCNVG
Q8JSZ3_CCHFV   LSEPRNIQQKLPPEIITLHPRIEEGFFDLMHVQKVLSASTVCKLQSCTHGVPGDLQVYHIGNLLKGDKVNGHLIHKIEPHFNTSWMSWDGCDLDYYCNMG
               :*:*:** .*** .::*::   :..*.*:*** :*:**.. *:******* .   :.  :  *:*.*: .*  :  ::*:.*:**:**:***:*****:*
Prim.cons.     2S2P2NI22KLP2222T22222222F2D2MHV22V2SA222C2LQSCTHG222222222222L2K2D222G2222222P22N2SW2SW2GCD2DYYCN2G

                     1410      1420      1430      1440      1450      1460      1470      1480      1490      1500
                        |         |         |         |         |         |         |         |         |         |
Q02004_DUGV    DWPTCTYTGVVTQKLREFLKLDQHRKRLHTTLSFSLKKNLSKRSHTSVRLEGKTVTRMEVKVTALIEVDGMELHSKTIRLSGIRLTGLKCSGCFSCTSGI
Q8JSZ3_CCHFV   DWPSCTYTGVTQHNHASFVNLLNIETDYTKNFHFHSKRVTAHGDTPQLDLKARP-TYGAGEITVLVEVADMELHTKKIEISGLKFASLACTGCYACSSGI
               ***:******. ::  .*::* : ..   ..: *  *:  :: . ..: *:.:. *    ::*.*:** .****:*.*.:**::::.* *:**::*:***
Prim.cons.     DWP2CTYTGV2222222F22L222222222222F22K222222222222L2222VT222222T2L2EV22MELH2K2I22SG22222L2C2GC22C2SGI

                     1510      1520      1530      1540      1550      1560      1570      1580      1590      1600
                        |         |         |         |         |         |         |         |         |         |
Q02004_DUGV    SCSVNAKLTSPDEFTLHLRSTSPNVVVAETSIIARKGPSATTSRFKVFSVRDTKKICFEVVEREYCKDCTPDELTTCTGVELEPTKDILLEHRGTIVQHQ
Q8JSZ3_CCHFV   SCKVRIHVDEPDELTVHVKSDDPDVVAASSSLMARKLEFGTDSTFKAFSAMPKTSLCFYIVEREHCKSCSEEDTKKCVNTKLEQPQSILIEHKGTIIGKQ
               **.*. :: .***:*:*::* .*:**.*.:*::***   .* * **.**.  ...:** :****:**.*: :: ..*...:** .:.**:**:***: :*
Prim.cons.     SC2V222222PDE2T2H22S22P2VV2A22S22ARK2222T2S2FK2FS2222222CF22VERE2CK2C2222222C2222LE2222IL2EH2GTI222Q

                     1610      1620      1630      1640      1650      1660      1670      1680      1690      1700
                        |         |         |         |         |         |         |         |         |         |
Q02004_DUGV    NDTCKSKIDCWSNSISSFASGIGDFFKHYIGSIAVGVLGTVLPFALLILFFIYGDKMLWPFKVFCRPCRRCCRKNEG---YNKLAEEEE--LRDIIRKFS
Q8JSZ3_CCHFV   NSTCTAKASCWLESVKSFFYGLKNMLSGIFGNVFMGIFLFLAPFILLILFFMFGWRILF--------CFKCCRRTRGLFKYRHLKDDEETGYRRIIEKLN
               *.**.:* .** :*:.**  *: :::.  :*.: :*::  : ** ******::* ::*:        * :***:..*   *.:* ::**   * **.*:.
Prim.cons.     N2TC22K22CW22S22SF22G222222222G2222G222222PF2LLILFF22G222L2PFKVFCRPC22CCR222GLFKY22L222EETG2R2II2K22

                     1710      1720      1730      1740
                        |         |         |         |
Q02004_DUGV    K---SGELINKDAKDKRTLARLFMSDNPKLKKEKKLSEIA
Q8JSZ3_CCHFV   NKKGKNKLLDGERLADRRIAELFSTKT----------HIG
               :   ..:*:: :   .* :*.** :..          .*.
Prim.cons.     2KKG222L22222222R22A2LF2222PKLKKEKKLS2I2
```

**Table S4: Sequence alignments and numbering of different *Peribunyaviridae* M polyproteins.** $Gn^{cyto}$ and NSm zinc fingers are highlighted in cyan and magenta, respectively. In grey, additional amino acids used in AlphFold predictions. Red type, green type, and blue type indicate identical amino acid residues, conserved substitution, and semi-conserved substitutions respectively.

```
                                    10        20        30        40        50        60        70        80        90       100
                                     |         |         |         |         |         |         |         |         |         |
Q806D6_9VIRU_AKAV           MIITILNVLTLVVTVTAMPPRNTNGGRCFYGGDMFRQINSTSPMSEICVRDDISLVKSIGYHKLAANREV-IESSMSYYRLYYVKNWFECNPVQDILGTF
A0A1I9WAL5_9VIRU_CEV        -------MIFMLILFAVAAASPVYQ-RCFQDGAIVKQNPSKEAVTEVCLKDDVSMIKTEARYT--KNSTG-VFSNNVAIRKWLVSDWHDCRPRKVVGGHI
F6JSE3_BUNLC_LACV           -------MIYILVLIAVTAASPVYQ-RCFQDGAIVKQNPSKEAVTEVCLKDDVSMIKTEARYI--RNATG-VFSNNVAIRKWLVSDWHDCRPKKIVGGHI
D7F1G7_9VIRU_BATV           ----------------------------------SRKESQAAISEFCLKDDVSTIKSEIIYE--KNDTG-LFAHNKVLRNWIIKDWKNCNPVPTAGGSI
Q6PN63_9VIRU_NRIV           -----MLLLLVLLIPCYVSASPVVT-RCFHGGQLIAEKKSQTAVSEFCLKDDVSTIKSEITYE--KNNTG-LFAHSKVLRNWIIKDWKNCNPVPTAGGSI
GP_BUNYW_BUNV               ----MRILILLLAVTQLAVSSPVIT-RCFHGGQLIAERKSQTSISEFCIKDDVSMLKSEIVYT--KNDTG-IFGHSKVFRHWTITDWKACNPVVTAGGSI
A0A0R7FK51_9VIRU_UMB        ------MMVFALLLISLAGLSSQATIRCFSGGSKIMERTSNVSLSELCLKDDVSQLKIIVDHT--KNDSG-LFSTASVWRKWIVSDWTTCNPKKMALGTI
A0A0R7FK55_9VIRU_WIT        ----MGFLNWLLLFFAMHTVRAIPSTRCFAGGQKMHELTSSKVTSEACLKDDVSQVKIHTNYF--KNETG-IYAIMTAYRKWTVADWHECRPKKTVSGSI
A0A2I4SI68_9VIRU_TCM        -----MELILFVTMIAAVLANPTPE-RCFTGGTKLLESKSPQAMTYVCLKDDISYVKSEVVSTDEKDKNGRTLFLNTVFRKIQIDDWQNCRPEKQVGGPI
Consensus                                             RCF gG    #  S    se ClkDD!S vKse      k#  g   s     Rkw ! #W  CnP k  gG i
Prim.cons.                  MIITMM6LILLLLLIA3AAASPV2T3RCF2GGQKI2E3KSQ2AVSE2CLKDDVSMIKSEI2YT22KNDTGR2FSHNKV4RKWIV2DW2NCNPVKT2GGSI

                                   110       120       130       140       150       160       170       180       190       200
                                     |         |         |         |         |         |         |         |         |         |
Q806D6_9VIRU_AKAV           MVFDVNHEGILAPKTYACRATCSISLERDTGNVVLESPALNHYTIHGTTIKNGWFKTKVSIDLDNTCEDLHITCGHKTLNVHACFRQHKSCIRYFKGSIL
A0A1I9WAL5_9VIRU_CEV        NVIEVGDDLTLHTESYVCSADCTIGVDKETAQVRLQTDTTNHFEIAGTTVKSGWFKSTTYISLDQTCEHLKVSCGPKSIQFHACFNQHMSCVRFLHRTIL
F6JSE3_BUNLC_LACV           NVIEVGDDLSLHTESYVCSADCTIGVDKETAQVRLQTDTTNHFEIAGTTVKSGWFKSTTYITLDQTCEHLKVSCGPKSIQFHACFNQHMSCVRFLHRTIL
D7F1G7_9VIRU_BATV           NVIEVNTDLSLTTKTYVCSRDCTITVDKEDAQIIFQTEKLNHFEVSGTTLSSGWFKTKASVTLDRTCEHIKVTCGKKILQFHACFKHHMSCIRFFHGTIL
Q6PN63_9VIRU_NRIV           NVIEVNTDLSLTTKTYVCSRDCTITVDKEDAQIIFQTEKLNHFEVSGTTLSSGWFKTKASVTLDRTCEHIKVTCGKKTLQFHACFKHHMSCIRFFHGTIL
GP_BUNYW_BUNV               NVIEVDKNLNLVTRNYVCTGDCTITVDRKNAQIIFQTDKLNHFEVTGTTISTGWFKSKASVTLDRTCEHIKVSCGKKTLQFHACFKQHMSCVRFLHRSIL
A0A0R7FK51_9VIRU_UMB        SVIDIDKDFILKSSSYICNKECTITVDKERAQVLLQSDGLNHFEVSGTTLSSGWFKLTATVTLDQTCEHVKVSCGQKSMQFHACFKNHMSCIRFLHRSIM
A0A0R7FK55_9VIRU_WIT        NVIEVDKSMMISSVVYTCDKDCTINIDKENAQIILHTEGINNFEVSGTTVIKAWFKTTATIPLQQTCEHIKIICGKKILQFHSCFKHHMSCIRYLQNSMV
A0A2I4SI68_9VIRU_TCM        MIISFDEHGSMESEDYVCKNDCEIKLDRDAGLINLETSTLNYYQISGTTKVSGWFKTRTSISLKHTCENIKIQCGEKSILLHACFKNHMECYQFLISHML
Consensus                    !i vd    l    YvC  dCtI  #r# aq! l t  lNh%e!sGTT  sgWFKt  s! Ld TCEh k! CG K lqfHaCFk HmsCir%l  sil
Prim.cons.                  NVIEVDKDLSL2T22YVCSADCTITVDKETAQIILQTD2LNHFEVSGTT2SSGWFKT2ASITLDQTCEHIKVSCGKKSLQFHACFKQHMSCIRFLHR2IL

                                   210       220       230       240       250       260       270       280       290       300
                                     |         |         |         |         |         |         |         |         |         |
Q806D6_9VIRU_AKAV           PEVMIESICTNAELILLCCFSAISCFVAIILTKTYLVYLLIPIFYPFVKLYGLLLQRFCKQCKNCLLPIHPFSPCPTTCICGMVYNSTEALKVHRKCLNC
A0A1I9WAL5_9VIRU_CEV        PGSIANSICQNVEIIILVTLALLIFILLSIISKTYICYLLMPIFIPIAYIYGLIYNKSCKKCKLCGLVYHPFTECGTHCVCGARYDTSDRMKLHRASGLC
F6JSE3_BUNLC_LACV           PGSIANSICQNIEIIILVTLALLIFILLSILSKTYICYLLMPVFIPIAYVYGVIYNKSCKKCKLCGLVYHPFTECGTHCVCGARYDTSDRMKLHRASGLC
D7F1G7_9VIRU_BATV           PGTMATSICQNIELIIIISLTLIIFILMVILTKTYICYLLMPLFMPIAYFYGWSYNKSCKKCPCCGLAYHPFTNCGSHCVCGLKFEASDRMRIHRESGLC
Q6PN63_9VIRU_NRIV           PGTMATSICQNIELIIIISLTLIIFILMIILTKTYICYLLMPLFMPIAYFYGWSYNKSCKKCSCCGLAYHPFTNCGSHCVCGLKFEASDRMRIHRESGLC
GP_BUNYW_BUNV               PGSMAISICQNIELIIITILALCIFIIMIILTKTYICYVLIPVFMPIAFAYGWAYNRSCKKCTCCGLAYHPFTNCGSYCVCGSKFETSDRMRMHRESGLC
A0A0R7FK51_9VIRU_UMB        PASFAVSICQNVELILMTTFTLIIFLLLSILTKTYICYILLPIFIPFAYAYGFIYDKTCKKCSLCGLAYHPFTKCGIHCVCGCRFENSDAMKTHREKGKC
A0A0R7FK55_9VIRU_WIT        PGKISNAICQNLELLIMTTITTLIFSILILLAKTYLCYLLLPLFIPIAYCYGFFYDRSCKKCKSCGLAYHPFTKCGEHCVCGGKFETSERMKIHRDSGLC
A0A2I4SI68_9VIRU_TCM        PKTFSSSVCANLEIIILTAFSIIAFLLLSILTKTYICYILLPFFIPCTYIYSKLYMKCCKICNNCGLRYHPFSPCGKICVCGSVYDSSERMKIHRQTGLC
Consensus                   P     s!CqN Eliilt f  iif ll iLtKTYicYlL P FiP ay Yg  y k CKkCk CgL yHPFt Cg  C!CG  %# s#r$k HR  glC
Prim.cons.                  PGSMANSICQNIELIIILTTLTLIIFILL2ILTKTYICYLLMP2FIPIAY3YGWIYNKSCKKCK2CGLAYHPFTNCG2HCVCG3KFETSDRMKIHRESGLC

                                                              BUNV NSm 1        10        20        30        40        50
                                                                       |         |         |         |         |         |
                                   310       320       330       340       350       360       370       380       390       400
                                     |         |         |         |         |         |         |         |         |         |
Q806D6_9VIRU_AKAV           TGYKTLTKTRYLCKKRLPNMVLATISTVLFFTFITPITA--------ECYNYTSLPADFQAAIDKNN----SCTREQLIMIILTILCIASTIMVISAVHL
A0A1I9WAL5_9VIRU_CEV        PGYKSLRAARVMCKSKGPASILSIITAVLILTFVTPINAMMVG-ESKETFELEELPDDMLEMANNLNTYYMLCVANYAVTWSLLIVALIIGLLFKKYQHR
F6JSE3_BUNLC_LACV           PGYKSLRAARVMCKSKGPASILSIITAVLVLTFVTPINSMVLG-ESKEVFELEELPDDMLEMALRINFYYFTCILNYAVSWSLVIIGLLTGLLFKRYQHK
D7F1G7_9VIRU_BATV           QGYKSLRVARLLCKSKGSSLIISILLSMLILSFVTPIEGTLTNYPESRKYDLEEIADVLEGFVVEKGIKEYVVFYTSIFGALFLLMALIVIIALNRVTEY
Q6PN63_9VIRU_NRIV           QGYKSLRVARLLCKSKGSSLVISILLSMLILSFVTPIEGTLTNYPESRKYDLEEIADVLEGFIVEKGIKEYVVFYTSIFGALLLLIALVITITLNKVTEY
GP_BUNYW_BUNV               QGFKSLRVARRLCKSKGSSLIISILLSVLILSFVTPIEGTLTNYPTDQKYTLDEIADVLQAKTHEDSTKYYIILYTSLFGAGLTIIFAGVALGLTIILEV
A0A0R7FK51_9VIRU_UMB        PGFKSMRTARVLCKAKGSAFTLSVLLATLALGFLTPVQGLILD-ETQASCTEDQLPESFQVMQSTISVFKFATIMNAGWTYCMFLITLALYVIYKKYAHK
A0A0R7FK55_9VIRU_WIT        QGYKSLRSARVLCKSKASALTLSVLLSILLLGFLTPVQGVPI----KTNFTIDELPEYSQDQSNQVTLLHKNVIYSLLVDLLLIILVFVSIICINHFVYS
A0A2I4SI68_9VIRU_TCM        PGYKYMMSARFMCKSKGWNMIVAILLSVLFFSFITPIGA------NSTCYTLDEFPDMYINSIKNMEFIKAYLFYWLAVNSLLLLIFITVIITIHYFEHI
Consensus                    G%Ks$r aR $CKskg    lsill vL l F TP! a          ytldelpd  q  i                   $ ii     i      h
Prim.cons.                  2GYKSLRVARVLCKSKGSALILSILLSVLILSFVTPIEGTLTNY232EKYTL2ELPD2LQ3MI2E2N3KYYV22Y2S2VGALLLIIAL3VII2L2KY3H2

                            60       70        80        90       100       110       120       130       140endNSm
                            |         |         |         |         |         |         |         |         |  |
                                   410       420       430       440       450       460       470       480       490       500
                                     |         |         |         |         |         |         |         |         |         |
Q806D6_9VIRU_AKAV           YIRIYYVFCAYCGMVHEKKGLILLDNFTSHCLTCICRDKS------IHRANNNCVTPMKYKTAKYTNLLCIILLIIIAVTPAFSA-------CIQEKEIE
A0A1I9WAL5_9VIRU_CEV        FLDIYAMYCEECDMYHDRSGIKWNGDFTNKCGHCTCGQYEDASGLLVHRKTYNCLIQYKSKWIMNFLVTYVILVLLKDSVVIVQAKGVDFSECIEAEKVR
F6JSE3_BUNLC_LACV           FLNIYAMYCEECDMYHDKSGLKRHGDFTNKCRQCTCGQYEDATGLMTHRKTYNCLVQYKAKWIMNFLIIYIFLVLIKDSAIIVQAAGTDFSSCLETESIN
D7F1G7_9VIRU_BATV           LTNINVLYCHECSMYHSKKNIKYIGDFTNKCGFCTCGELEDQEGLKIHKVSRKCIYKYQLTWSKVIMMILVCLLIAQNTILIVAAS----DDCWTKKSLE
Q6PN63_9VIRU_NRIV           LTNINVLYCYECSMYHSKKNIKYIGDFTNKCGFCTCGELEDQEGLKIHKVSRKCIYKYQLTWSKIIMTILVCLLIAQNTILIVAAS----DDCWTKKSLE
GP_BUNYW_BUNV               LTKINVIFCNECNMYHSKKSIKYVGDFTNKCGFCTCGLLEDPEGVVVHKAKKSCTYSYQINWVRGIMIFVAFLFVIQNTIIMVAAE----EDCWKNEELK
A0A0R7FK51_9VIRU_UMB        LLNYYVIHCKECDMLHDKSGLKYNGDFTNKCGFCTCGQVEDAMGLTVHQIRNHCSANYRTRMVLNWMAVLLATIIKDSIMLAAAAGTEFETCVKKVEMT
A0A0R7FK55_9VIRU_WIT        IANIFAIYCNTCDMYHSRSNLKYFGDFTNRCGSCTCGQFEDMQGVLIHKRSSRCLTKYKIMTLKHVILWIIITLVVKDFAMVAQAK--TLKECLQIKELE
A0A2I4SI68_9VIRU_TCM        YLRPYIFKCEDCKMFHSKHRLVIEEFGTSRCGSCTCGCPDDPPMVHFHQTSRQCVLKYIVKILKTYSICIILYCFISSTTIGAIAT--EVGVC-DDDPID
Consensus                      iy   C  C M H k glk  gdfTn Cg CtCg  ed  g   H     C   yk k  k      i l  i        A       C    ei
Prim.cons.                  LLNIYVIYCEECDMYHSK2GLKY2GDFTNKCGFCTCGQLEDAEGL2IHK3SRNCLYKYK2KW4KNIMIILIILLIIKDTIIIVAA3GT2F3DCW4KKELE

                                   510       520       530       540       550       560       570       580       590       600
                                     |         |         |         |         |         |         |         |         |         |
Q806D6_9VIRU_AKAV           T-IDDAAMCIGLYQNITQPKEYNTFVKELSSTLSSHEIEF-LLPAVKPSFDYLITKSSTSKDLHSATIHEFIAANLYPNNFKKHLASAGPGSIQWRTYIQ
A0A1I9WAL5_9VIRU_CEV        WNCTGPFVNLGNCEKKQKKEPYSNIAAQLKGLGAISILDVPIIASIPEDIAGALRYIEDQRSYHSQLTVEYAMLTRYCDYYTQFSDNSGYSQTTWRVYLR
F6JSE3_BUNLC_LACV           WNCTGPFLNLGNCQKQQKKEPYANIAAQLKGLKAISVLDIPMITEIPDDIASALRYIEEREAFHVQLTAEYAMLTRYCDYYTQFSDNSGYSQTTWRVYLR
D7F1G7_9VIRU_BATV           MECIGPLQQVDTCDDKTSRS-YSGEAKKLVRASKISDLDAAQVGLLGSTIESAIASIRRQRTYSTMHLLESVFLGKHCDYYKVFEHNSGYSQAKWRLTAK
Q6PN63_9VIRU_NRIV           MECIGPLQQVDTCDDKASRS-YSGEARKLVRASKISDLDAAQVGLLGSTIESAIASIRKQRTYPTMHLLESVFLGKHCDYYKTFEHNSGYSQAKWRLTAK
GP_BUNYW_BUNV               EDCVGPLIAPKDCTDKDHKT-YLSEASLLATAKKITQVDAENVEILGKTMESAIRVIERQKTYHRMHLLEAVFLNKHCDYYKMFEHNSGYSQVKWRMMIK
A0A0R7FK51_9VIRU_UMB        PECTGPFMEVGICDHQQKSLKYHVVANMLLSKGTISKLDVDIIDALPDKIAGDLQMIQSSSDFHHQQMLEYVFLLRHCDYYSSFVSDSSYSQILWQTAAK
A0A0R7FK55_9VIRU_WIT        KDCTGPFFDIKECSTTTMQKSYGDIANWLKTKEVISELDKKAASELSQDVTRSLQKIETTLGLHQKILFEILFLSRNCDYYKSFDNVYSTHQLQWQTLAK
A0A2I4SI68_9VIRU_TCM        TECWGLNIEKDIISNRNDIN-----GTLRKMFPDMTEEELNIAHTDYATYPDFLIEARSYDALRPKQILEAVYFSRNSFHHITGADQRYY--ISWKLRTR
Consensus                     c gp    g c        y   a  l     i e #              l  i     lh     E   l r cdyyk f    gy qi Wr
Prim.cons.                  3ECTGPF33VG3CDDK2KK3PYS22A33LK4A3KISELDA2I2G3L22TI2SALR3IE3QRTYH22HLLEYVFL4RHCDYYK3FE2NSGYSQIKWR2YAK

                 610       620       630       640       650       660       670       680       690       700
                                     |         |         |         |         |         |         |         |         |         |
Q806D6_9VIRU_AKAV           NNNLHLCNDHVVKMICRCVIKQEECSS-TKVDDGEQIAQYYKRNKEFYKADLEILYTVISRAIPGLVGNLLRQVLKSQKYGESLHVLNKIKKDVSKN---
A0A1I9WAL5_9VIRU_CEV        SHDFEACVLYPNQHFCRCVKHGDKCSS-SNWDFASEMKNYYSGKQSKFDKDLNLALTSLHHAFRGTSSSYIAGLLSSKANDELVQYTSKIREKFPGNALL
```

```
F6JSE3_BUNLC_LACV      SHDFEACIMYPNQHFCRCVKNGEKCSS-SNWDFANGMKDYYSGKQAKFDKDLNLALTALHHAFRGTSSAYIATMLSKKSNDDLIAYTNKIKTKFPGNALL
D7F1G7_9VIRU_BATV      THHFDICSRHSTHHFCRCISDGTKCQN-GDWDFAGEMNSTYQSKKDFFEHDFKLFCTLVENAFPGTTESLFYEMLSKKNTTGITKLLDKLTKKFGNNNMF
Q6PN63_9VIRU_NRIV      THHFDICSRHSTHHFCRCISDGTKCQN-GDLDFAGEMNSTYQSKKDFFEHDFKLFCTLVENAFPGTTESLFYEMLSKKNTTGIKQLLDKLTRKFGNNNMF
GP_BUNYW_BUNV          TQHFDICALQANSPFCAQCIADNSCAQ-GSWEFDTHMNSTYSSKVDNFKHDFSLFLRIFEAAFPGTAYVHLLTNIKEKKPYQAVSMIEKIKKKFPNNKLL
A0A0R7FK51_9VIRU_UMB   LNETNLCKHESSKKSCKCLKSNANCGE-SE-DINTAAQTYYKNKESELGQDLTTIMYILKKMIPGSGYSYLLNATKHNKLELFKAYMTNISSKYPNNKKL
A0A0R7FK55_9VIRU_WIT   NGDFDYCKTKGTEALCKCI-SGAACQS-II-DKKDDIDTHYKSKDQQRKKDIETVMSITRYMFPGTGHSYIANQTVTKQYTKVVEYFTSFAEKHATNLRL
A0A2I4SI68_9VIRU_TCM   VLELNTCKLVAKPYPCKCISIEEECTEFALKKFRSANLDNYLTDQTKLNWDRLKLQDAMYEFMPPTVVHKYISIVESKNKTMIDGFSHDIYQMYTN----
Consensus                    C        C C    e C s    df      yY  k      Dl         a pgt          sk          ki  k   n
Prim.cons.             TH2FDICKLHS23HFCRCI22GEKC2SF22WDFA3EMNSYY2SK2D2FK2DL3LFLTILE2AFPGT43SY2A3MLS2K23T2IV2YL3KIKKKFPNN3LL

                              710       720       730       740       750       760       770       780       790       800
                                |         |         |         |         |         |         |         |         |         |
Q806D6_9VIRU_AKAV      -------NQLNGIVEFLIHINSQNITEEVRELRIRPDLSIRGSKFTDKN----PGIPDIKECQTPLFITCTGKRFR-SLMKQYIAC-SNGGVKLYQRPNK
A0A1I9WAL5_9VIRU_CEV   KAVIDYIAYMKGLTEMSNFKKDEFWDELVYVPDPTKPPKLSRSSGTVYDFKSATSNLGEKNCKDVKGVVCLSPRSG-VTYDSIIACGEGSSPNIYRIPKT
F6JSE3_BUNLC_LACV      KAIIDYIAYMKGLPGMANFKHDEFWDELLYKPTPTKAPNLARGRESSYNFKLAISSKSIKTCKNVKDVACLSPRSG-AIYASIIACGEPNGPSVYRKPLN
D7F1G7_9VIRU_BATV      VGIWKFGQYLMSLPYVNETSLSPIQLAKVLEVTDQHHRSVSGRQESLAS---ATPGSKSSECDHAKKVSCISPRFG-APMEGLMACGNSPNYKIYKTPAK
Q6PN63_9VIRU_NRIV      VGIWKFGQYLMSLPYINETSLTPAQISKILEVTDQHHRSISGRQESLAS---ATPGSKSKECGHAKKVSCISPRFS-VPMEGLMACGDSPNYKIYKTPTK
GP_BUNYW_BUNV          IGYLDFGKYLLGLSHASTYELQQRQLDKLYQPTELTRSG--GQQTSLAN---SVVGQATKECKKYKDVSCLSPRFG-IPLEDLISCCDQPNYNIYKKPKK
A0A0R7FK51_9VIRU_UMB   KDFMAITNAILANLSALSHSSEESIPEYKQIDFSSSVGETTDMLEANFN---QTISTDKITCENPKVMICISPRTK-ISSGELYVCQRSGKYYIIDTGRF
A0A0R7FK55_9VIRU_WIT   KAFSILISGLLKTTTSLE--PPEQLASFVPKLARPTIPGFTDLIDSNFD---QSAQNDGRRCKSPSVIVCASPRTG-YTSDEMYIC-RDTKYYIIDTKGL
A0A2I4SI68_9VIRU_TCM   --YPYILNFMYLLMNITSRSQDFSISALLQNQTHATTQILNRGLVTIES----LELGYRKTCKNPKRYFCKSERVGLVATKPLLVCDRDTSNGETKRTLH
Consensus                     n l  l           i e                  t  n          k Ck pk   C spR g         C        y  p
Prim.cons.             K2I2DF2NYLLGLP23LE3SLDE222E2VYEPT22THPSLSG3QESLANFK2AT2GSD2KECKNPK3VSC2SPRFGLVPME3LIACGRSPNYKIYKTP2K

                              810       820       830       840       850       860       870       880       890       900
                                |         |         |         |         |         |         |         |         |         |
Q806D6_9VIRU_AKAV      PLA-----LVGNKLCIGDKYCMIAFDPMVIDE--NIQKLDCYSLAATDQSDGMLKPE-RSIRLLKTGECKIAGA--LSRIAVSINQKNYKYSTIVHKKSG
A0A1I9WAL5_9VIRU_CEV   GVYQ--SNSEPSNYCVSDSHCLEDFEVINQEELDAIKKSRCWEVDYLYANPSKQSDGIRSCRMRDAGHCNVTTNR-WPIIQCDDNKYYYSELQRDYDKEQ
F6JSE3_BUNLC_LACV      GVFQ--SSTDQSMYCLLDSHCLEEFEAISQEELDAVKKSKCWEIEYSDIKPLQESDGTKSCRMKDSGNCNVATNR-WPVIQCENDKFYYSELQKDYDKTQ
D7F1G7_9VIRU_BATV      -LYK--SNNKGEVWCSGDVHCFQELTPASQESVDRIKQMTCFLTE-PEVNDDVFSIAIAMCKVQDKGVCTINEDR-WNIIKCDSGLIYYTDQREGQDTGN
Q6PN63_9VIRU_NRIV      -LYK--SNNKGEVWCSGDVHCSQELSPASQESVDRIKQITCFLTE-PEVSDDVFSIAISTCKVQDKGVCTVNEDR-WNVIKCDSGLIYYTDQRDGQDTGN
GP_BUNYW_BUNV          -VYK--AHDKEETWCINDQHCLVDFVPAEADTVEKLKPMKCWLVD-PGKNDDVYSIAIKTCRVVDKGVCTVNSQK-WNIIKCDSGPLYYSDHIPGEDTGN
A0A0R7FK51_9VIRU_UMB   KLYK--LIGAGNLYCAADAQCRYEFRILTTGEMQATFKDN-CKASVHTGTPGYQNDAVTHCKVLASGKCKYNNQTLIDITLCANNRYYPEAKGRI--FGV
A0A0R7FK55_9VIRU_WIT   QLYK--LDNSAGTYCVGDKYCNLKYRVLTPEESLALRKDNNCKHKEYTEPKNYLTERLTICRQKATGKC--GEEPTFRVSLCENGFVYPTAAKASPKDSA
A0A2I4SI68_9VIRU_TCM   RWSQDLVEVDQSIICRKDYTCSSYFDNSTDIELQTITKKKQRCNEIAKVYDPVYNQAISKCKMVEKGFCNFNGNP-KQIVKCSDNTLVEQIATGNYLNDM
Consensus               l            C  D  C   %      e   i k  c         d         cr    G C         ii c  n  yy
Prim.cons.             GLYKDLSNNKG22YC3GD3HCL2EF3PA2QEE2DAIKK3KCWL2EYP3VNDDV4S2AI4SCRV4DKGVC2VNENR2WNIIKCD2223YY2D33RGYD2GN

                              910       920       930       940       950       960       970       980       990      1000
                                |         |         |         |         |         |         |         |         |         |
Q806D6_9VIRU_AKAV      LVDEYCLSPNCDLDCYPYYPANLVDCSWSESTHSTLNQKVISHTDIESFISSVKLSLHNDLIQHHFRPLSNMPHVKPNFKSINVQGTISGGKIQDSYITF
A0A1I9WAL5_9VIRU_CEV   DIGHFCLSPRCTTVRFPIHPKHVSNCDWQVSHSTVDKVDVHNLEDIEQYRKAITQKLQNSLSIFKYARTKNLPHIKPIYKYITIEGTETAEGIESAYIES
F6JSE3_BUNLC_LACV      DIGHYCLSPGCTTVRYPINPKHISNCNWQVSRSSIAKIDVHNIEDIEQYKKAITQKLQTSLSLFKYAKTKNLPHIKPIYKYITIEGTETAEGIESAYIES
D7F1G7_9VIRU_BATV      DFGEYCLSHSCRIERFPINPAIISDCLWEYHSRKSKYITSLDLESLEEFKRAISEKLSHTLVVYNFKPTANLPHIKPVYKYITVQGVENSDGVDSAYITA
Q6PN63_9VIRU_NRIV      DFGEYCLSHSCRIERFPINPAIIRDCLWEYHSRKSKYITSLDLENLEEFKRAISEKLSHTLIVYNFKPTANLPHIKPVYKYITVQGVENSDGVDSAYIAA
GP_BUNYW_BUNV          DIGHYCVSAGCKTDRYPINPDVVTDCVWEFTSRKSQYIGKISMQSLEDYEKALTDRLTHTLETYSFAPLENLPHIKPVYKYITAQGVENSDGIEGAFITA
A0A0R7FK51_9VIRU_UMB   DPEDLCFDSACTKQQTPYNEELISACVWENPTLRPIRPRTEVSTTFQEYKDQLLKKINTDLVLHKFVRTQNLPYFIPQFKYITIQGTATTDGITSSFIFF
A0A0R7FK55_9VIRU_WIT   FPNDRCFDTTCKYGYYPYNMIQYSKCVWDNIKITSSRIKIASYENFQEYKDQLLKKITSDLTIGKFNLVENMPYFIPKRKYLTIKGVTTADGIDGSFVEF
A0A2I4SI68_9VIRU_TCM   KMDQVCFKTDCGDVK-RIHPDSLTNCTINVPRQLPVHIDRVDTNDFKIYKAHLEEDFLTTLSKFYFMPTKGLPHIVPDFRPIYLRGTETTDGLESSYFEI
Consensus                   C s  C    ypi p     C w         i      d e %k  l   l   L    % pt n$PhikP fkyit qGtet dgi ss%i
Prim.cons.             DIG2YCLSP2CTTVRYPINPA2ISDCVWEVSSRKSKYIDV32LED3EEYKKA2TEKL32TLS32KFAPTKNLPHIKPVYKYITIQGTET2DGIESAYIE2

                             1010      1020      1030      1040      1050      1060      1070      1080      1090      1100
                                |         |         |         |         |         |         |         |         |         |
Q806D6_9VIRU_AKAV      SIPLMTGLSQGFTLQDHKGNTLFDIIAYVKSARVIATYNHEYKTGPTVSINVQHNEQCTGSCPSSIPKKDN-WLTFSREHTSTWGCEEWGCLAIGTGCVY
A0A1I9WAL5_9VIRU_CEV   EIPALAGTSIGFKINTKEGKHLLDVIAYVKQASYSSLYNKMYVTGPTVGINTKHDELCTGPCPVNVPHSDG-WLTFARERTSSWGCEEFGCLAISDGCVF
F6JSE3_BUNLC_LACV      EVPALAGTSIGFKINSKEGKHLLDVIAYVKSASYSSVYTKLYSTGPTSGINTKHDELCTGPCPANINHQVG-WLTFAKERTSSWGCEEFGCLAVSDGCVF
D7F1G7_9VIRU_BATV      SMPALSGTSIGYNIMSRDNFPLFDIIIFIKSAIIKATYNHIYDTGPTIGINVMHDEHCTGQCPTDIPHKEN-WITFAQERTSRWGCEEFGCLAVNTGCVF
Q6PN63_9VIRU_NRIV      SMPALSGTSIGYNIMSRDNFPLFDIIIFIKSAIIKATYNHIYDTGPTIGINVMHDEHCTGQCPTDIPHKEN-WITFAQERTSRWGCEEFGCLAVNTGCVF
GP_BUNYW_BUNV          SIPAAGGTSIGYNVRSKDGFPLLDLIVFVKSAVIKSTYNHIYDTGPTISINTKHDEHCTGQCPSNIEHEAN-WLTFSQERTSRWGCEEFGCLAVNTGCVF
A0A0R7FK51_9VIRU_UMB   EIPALTGASVGYKVVNNEGQELIDMVVYIKTSKTSAKYKRVYYTGPTIAINTKHQEHCTGTCPERIEHDDG-WATFSKERTSNWGCEEFGCLAVETGCLF
A0A0R7FK55_9VIRU_WIT   EIPSLTGVSAGYTVTTKEGQELFDFIVYIKNSATSATYSYAYSTGPTIGINNKHTEVCTGKCPEKIPHEAG-WATFSRERTSTWGCEEFGCLAINEGCLY
A0A2I4SI68_9VIRU_TCM   EMIALSGKATGIKLYASNDLYLFDVVIYVQAANVTSTYVPIYRTGPTLTFNVKHEEICTGLCPSDLPRADNTWMTFSKESTSTWGCEEFGCLAIGEGCVY
Consensus              eipa$ G s G       g  LfD ! %!k a   atY   Y TGPT  iNvkH E CTG CPs iph dn W TFs ErTStWGCEEfGCLA! tGCv%
Prim.cons.             EIPAL2GTSIGYKI2SKEGFPLFD2I3YVKSA2ISATYNHIYDTGPTIGIN2KHDEHCTGQCPS2IPHKDNTWLTFS3ERTS2WGCEEFGCLAVNTGCVF

                             1110      1120      1130      1140      1150      1160      1170      1180      1190      1200
                                |         |         |         |         |         |         |         |         |         |
Q806D6_9VIRU_AKAV      GSCQDVIREEATVISRVNNEQLEVEFCVSEPTSTMCNTINVLEPVLGEHMQFEVHSVQTNLLPEVALIKNRRVYKGSINKKGVFNPQCGSVQSFDGKLYG
A0A1I9WAL5_9VIRU_CEV   GSCQDLIKEELAVYRKETEEATNVELCLTFSDKTYCTELNPITPIITNLFEVQFKTVETYSLPRVVAVQNHEIRIGQINDLGIYSKGCGNVQKVNNTIYG
F6JSE3_BUNLC_LACV      GSCQDIIKEELSVYRKETEEVTDVELCLTFSDKTYCTNLNPVTPIITDLFEVQFKTVETYSLPRIVAVQNHEIKIGQINDLGVYSKGCGNVQKVNGTVYG
D7F1G7_9VIRU_BATV      GSCQDIIRPETKVYRKAVEESVLLTVCITYPGKTFCTEINAIEPKITDELELQFKTVDTKTLPNILAVQNHKLYSGQINDLGSFSQGCGNIQKTNSSIIG
Q6PN63_9VIRU_NRIV      GSCQDIIRPETKVYRKAVEESVLLTVCITYPGKTFCTEINAIEPKITDELELQFKTVDTKTLPNILAVQNHKLYSGQINDLGSFSQGCGNIQRTNSSIIG
GP_BUNYW_BUNV          GSCQDVIRPETKVYRKAVDEVVILTVCITYPGHTFCTEINAIEPKITEEIELQFKTVDTKTLPYIVAVNNHKLYSGQINDLGTFGQMCGNVQKTNSSILG
A0A0R7FK51_9VIRU_UMB   GSCQDVIRRELDVFSKLEEDKILTELCITLSHDTYCVTIDSTTPTINDYFEIQYKTVDVQTLPKMIGVTHHKLLRGQINELGSFGKFCGNIQVVGNQTYG
A0A0R7FK55_9VIRU_WIT   GSCQDVVKPELDIYSKAGADTTKTNVCISLNHKTYCQEIDALNPIITENIEAQFKTVESKNLPRLIAIKNHKLYTGQINELGSFGKYCGNLQVTNNQTLG
A0A2I4SI68_9VIRU_TCM   GQCKNVIKPESEVWEKQSEERITLNLCITTSYEYYCKEIESTVPLITNKIEAQFNTIESFKHPERILIRNHIAFQGQINGLREYNSYCGNVQIVGSTTMG
Consensus              GsCq#!!r E  !  k  e#    e Cit    tyC ei#   P it#  # #fkt!#t  lP    ! nh  y GqIN lg %   CGnvQ      yG
Prim.cons.             GSCQDVIRPELKVYRKAVEE22LLE2CITY2GKTYCTEINAIEP2ITDE2ELQFKTV2TKTLPRI2AVQNHKLYSGQINDLGSFSKGCGNVQK2NS2IYG

                             1210      1220      1230      1240      1250      1260      1270      1280      1290      1300
                                |         |         |         |         |         |         |         |         |         |
Q806D6_9VIRU_AKAV      IGNPKFDYICHALSRKDIVVRKCYENHYYSCATLKE-AVEIKPNITNSKTMLYNDNALLGSASVKIMLGDLIYQQTSVQEKDIRGHATCGGCTDCFNDVA
A0A1I9WAL5_9VIRU_CEV   NGVPKFDYLCHLASRKEVIVRKCFDNDFQACKFLQS-PPSYRLEEDDGSITVIDYKKILGTIKMKAILGDVKYKTFA-ENIDITAEGVCTGCVNCFENIH
F6JSE3_BUNLC_LACV      NGVPRFDYLCHLASRKEVIVRKCFDNDYQACKFLQS-PASYRLEEDSGTVTVIDYKKILGTIKMKAILGDVKYKTFA-DNVDITAEGSCTGCINCFENIH
D7F1G7_9VIRU_BATV      TGTAKFDYVCHGASRKDIIVRRCYNNNYESCKLLKE-EQSLMFADNHETIDVANVRHLLGDLQFKLMLGDLRYKSFA-ENPDLEIEAKCVGCPSCFTSYS
Q6PN63_9VIRU_NRIV      TGTAKFDYVCHGASRKDIIVRRCYNNNYESCKLLKE-EQSLMFADNHETIDVANVRHLLGDLQFKLMLGDLRYKSFA-ENPDLEIEAKCVGCPSCFTSYS
GP_BUNYW_BUNV          TGTPKFDYTCHGASRKDIIVRRCYNNNFDSCKLLKE-ETQLIFNDDHDTITVYNTNHLIGELAIKLILGDIQYKLFT-ETLDLQIDAKCVGCPDCFESYS
A0A0R7FK51_9VIRU_UMB   QAAVKFDYICHAAQRKDVVIRKCLENNFQSCKLLTH-DTDLILEESETSVEVINNKRITGQIALKVLLGDFNYKQYT-ANVEADIEANCVGCTNCIKGIT
A0A0R7FK55_9VIRU_WIT   LADVKFDYTCHAAQRKDILIRKCLENSYQSCKLLKE-ESDYMLIEEATHVTISDNKKINGNLAIKAIFGDFNYKSYT-KDLDFEAEISCVGCFSCLHGIV
A0A2I4SI68_9VIRU_TCM   MGIPKMDYKCHAASRKDVIVRHCYDNNYLSCLELKPLPKDMIEYETDKLLTLNTHNKITGTLKLKIKLGDIQYKLFQ-EDTKLDITGECVGCIDCIDGIT
Consensus               g pKfDY CHaasRK#!i!RkCy#N % sCk Lke        e     t  n n i G    K  lGD  Yk f   # d    a CvGC dCf  i
Prim.cons.             TGTPKFDY4CHAASRKDIIVRKCY3NNYQSCKLLKELE2SL2LEEDH3TITVIN3KKILGTL24K2ILGDL4YKSFAVEN3DLEIEAKCVGCP3CFE2IS

                             1310      1320      1330      1340      1350      1360      1370      1380      1390      1400
                                |         |         |         |         |         |         |         |         |         |
Q806D6_9VIRU_AKAV      CKISMTSNGVYQCPIVSSCDSYINNVYINEGTNDVNLKFRCLKA----EIKISICGKEIPVKSEIIKDTKKLDWASADQTSYIKEFDKKCATWLCRAYNE
A0A1I9WAL5_9VIRU_CEV   CELTIHTTIEASCPVTSPCTLFHDRILITPDEHKYAMKVICTEK-QTGTLPFKICNSKVDATLTLVEAKPILELASVDQTAYIREKDERCKTWMCRVRDE
F6JSE3_BUNLC_LACV      CELTLHTTIEASCPIKSSCTVFHDRILVTPNEHKYALKVVCTEK-PGNTLTIKVCNTKIEASMALVDAKPIIELAPVDQTAYIREKDERCKTWMCRVRDE
D7F1G7_9VIRU_BATV      CSFKIVSNIDTVCSIEGPCTTFHNRLMITSTKQDYGIKMSCKTK-PGLTEEFQICKKTYTVLFATVEKNDKIEISTGDQTSFIQERDDRCKTWLCRVRDE
Q6PN63_9VIRU_NRIV      CSFKIASNIDTVCSIEGPCTTFHNRLMITSTKQDYGIKMSCKTK-PKETEEFQICKKTYTVLFTTVEKNDKIEISTGDQTSFIQERDDRCKTWLCRVRDE
GP_BUNYW_BUNV          CNFQIVSNIDTICSLEGPCDTFHNRISIKAMQQNYAVKLSCQKD-PRPSGTFKICNREYTVVFHTVAKDDKIEINVGDQTSFIKEKDDRCKTWLCRVRDE
A0A0R7FK51_9VIRU_UMB   CEIKVKTEIEASCKVEPPCPSYTNRIIIKPGRDTHTLLMKCDKKITSGELTIKICNMQIQAHLSTIESNDQLELSTGDQSTYIHEEDLRCSTWICKVRDE
```

```
A0A0R7FK55_9VIRU_WIT    CEANLKTTVETSCEVTSICSLYTNRLILLPSQEKYSLMIRCDRQIAENELDISICGKKIESHITITNTNDKLELSTGEQSTYIHEEDLRCTTWLCKFQEE
A0A2I4SI68_9VIRU_TCM    CRLEIPTSTQSTCTIKSNCETFINRIVISPNDKNYHLKFQCRTK--TDYVSLDICGNKKDISLKLSSVKPQLEVLPQAESAYVKERDDKCSTWLCKVKDQ
Consensus               C     t     C ! s C  % #ri i p    y lk  C  k       i ICg  i           kl#    d#t %!kE D rC TWlCrv ##
Prim.cons.              CE2KI3TNIETSC2IESPCTTFHNRI3ITP33Q2YALKMSC32KIP22TLT2KICNKKITV32TTVEKNDKLELSTGDQTSYIKE2DDRCKTWLCRVRDE

                              1410      1420      1430      1440      1450      1460      1470      1480
                                 |         |         |         |         |         |         |         |
Q806D6_9VIRU_AKAV       GIGFVLEPLWNELSLWGKYVLLALAIIISVLILVKVIRPLARYVVAVLKENDKVYKLENKLK--------------------------
A0A1I9WAL5_9VIRU_CEV    GIQVILEPFKNLFGSYIGIFYTSILIVITIFIIIYVILPICFKLKDTLQKHEDAYKREMKIR--------------------------
F6JSE3_BUNLC_LACV       GLQVILEPFKNLFGSYIGIFYTFIISIVTLLVIIYILLPICFKLRDTLRKHEDAYKREMKIR--------------------------
D7F1G7_9VIRU_BATV       GISVIFEPIRAFFGSYFSIAFYIVIGIIVLFLVIYIFLPMMFKLRDVLKRNEYLYLQEIKHK--------------------------
Q6PN63_9VIRU_NRIV       GISVIFEPIRAFFGSYFSIAFYVVISIIVLFLAIYIFLPMVFKLRDVLKRNEYLYLQEIKHK--------------------------
GP_BUNYW_BUNV           GISVIFEPIKAFFGSYFSIFFYIIVVVVVGFLIIYIFMPMFMKLKEVLKANEKLYLQEIKQK--------------------------
A0A0R7FK51_9VIRU_UMB    GTSFIFKPLTDWLGSFTKPILIAGCIIIVTIILIYILMPMCARLRDLLKKNEYEYLQDTKVIVKPKTAANSIGLRSKKINIQSSKAVD
A0A0R7FK55_9VIRU_WIT    GFNFILGPIYNWLGKFTWPVIAIIIIIFLIFIGVYVFMPMCMKLRDILKKNEYEHLQEIKTDMNNIPMIVKYKIKPFA----------
A0A2I4SI68_9VIRU_TCM    GIGFLFSPFTGFFGEVWHYVVLFLAVMITFFIVVYILIPCLFKCRDILRKQQAEFIVERKLR--------------------------
Consensus               Gi fileP  n fg     v     iii  fi !y!  P   klrd Lkkn#  y  # K
Prim.cons.              GISVIFEPIKNFFGSYFSI2FYIIIIIIVLFIIIYIFLPMCFKLRDVLKKNEY2YLQEIK3K2222222222222222INIQSSKAVD
```

**Table S5: Sequence alignments of different *Phenuiviridae* M polyproteins**. NSm is highlighted in grey. Red type/ asterisk (*), green type/colon (:), and blue type/dot (.) indicate identical amino acid residues, conserved substitution, and semi-conserved substitutions respectively.

```
                           10        20        30        40        50        60        70        80        90       100
                            |         |         |         |         |         |         |         |         |         |
GP_RVFV            -MYVLLTILISVLVCEAVIRVSL-----SSTREETCFGDSTNPEMIEGAWD---------SLREE-EMPEELSCS------------------------
A0A143Q355_TOSV    -MFIAKLLLISCLVQQAYLKYTIISEIGNGTRTSFCYSSRTSLGSLLEAWDNITESIGLTSGTCKVGSLENRDCKRDIMSNYVAEVHSNLHGLGAMMHNY
I1SV55_9VIRU_AMTV  ----------------------------------------------------------------------------------------------------
GP_PTPV            MIFTILNVLTRAMLVMSMYSLTTWD---STSRNDMCFSNDSPLEGLVYYWET-------HSKRHDYKKQESQRCR-----------------------V

Prim.cons.         MMF33L33LIS3LV33A3333T322EIGS3TR333CFS33T3LE3L33AWD2ITESIGL2S3R332333E333C3RDIMSNYVAEVHSNLHGLGAMMHN2

                          110       120       130       140       150       160       170       180       190       200
                            |         |         |         |         |         |         |         |         |         |
GP_RVFV            -------------ISGIRE-----VKTSSQEL----------------YRALKAIIAAD-------GL--NNI--------TCHGK-----------DPE
A0A143Q355_TOSV    GDDKRVEVLNDVIPDELNPGRHNCSDLDRKPFWTEFIDVAKPTNKIAPMSVIPPPIVKPIPV--AESTIKKSIP-FDKLLSEAEKELNQT-KIRLETERR
I1SV55_9VIRU       --------------------------MHQITVVSGPTEVSTCFGSLHPFQSLKPVMANALGV--LEGKMFCSIG-GRSLEEDDEGVLAVV-QGLQDSQQE
GP_PTPV            GDSDKKMITNVTIISLISE-----IQKSISELSLSCVNDDNSTGQVLTFNGLEDTIRGDYIVDCVTGLYQSDIGVGVGLGRTHHGHQQMKNKAVVIDEKE
                                                  .                   :   :          .    .*          .             : .
Prim.cons.         GD2222222N22IIS3I3EGRHNC334S44EL333333V333TG333PF44LKP4IA4D33VDC3EGL334SIGVG33L33T42G4L333NK33333E4E

                          210       220       230       240       250       260       270       280       290       300
                            |         |         |         |         |         |         |         |         |         |
GP_RVFV            DKISLIKGPPH----------------KKRVGIVRCER---RRDAKQIGRETMAG-------------------------------IAMTV--LPALAVFA
A0A143Q355_TOSV    SNAEEAAKERARMREIEDKLRKDWELERERKDAVADDLA--KEIARRREIERKAHELDKMLKKKEREDLVARERVSEDMRNHVRKPNTTIAPAPTIITTL
I1SV55_9VIRU       EEISCHGTPKGKLVKYKRHPRK----ASGKKGAVDCVSG--KELKPQKKRGDLAH--------------ISEHFQADEVRSSSVNISTTL---KTILVAA
GP_PTPV            RMISLLETQQSENDIKMQVLMSEIEQLKNQLSKKRNERGQEKRDAEKVMSDLMARNSDLRKHNDILTAEISQMKNKNTIQRNKNTVSTTV--VPAILSVA
                      .                         : .         :.   :      *                                  *:    ::
Prim.cons.         44ISL44TP4433333333LRK22E23K4RKGAVRCERGQEK2DA4Q44RE4MAH22D22222222222IS33333333R33333ISTTVAP3P2ILV4A

                          310       320       330       340       350       360       370       380       390       400
                            |         |         |         |         |         |         |         |         |         |
GP_RVFV            L-----APVVFAED--------PHLRNRPGKGHNYIDGMTQEDATCKPVTYAGACSSFDVLLEKGKFPLFQSYAHHRTLLEAVHDTIIAKADPPS-CDLQ
A0A143Q355_TOSV    LMA---SAVMAGPLEN---RETNHLLNRPGNGAYTLSDFAES--TCT-LAYGSECKSWEHQLDELSFPFFHSNLDKYSMLEAATETIPILNKSSAVCTIS
I1SV55_9VIRU       LLT---SALVNG---------DPHIDNRPGKGDYA-RDASRSGNNCSPLKYGAACKAWTYQKDVSTYPFFNTFPYKYSLIEALSEGFIQMDEQNT-CKLT
GP_PTPV            LLSSSVAPIIAAPPDSPMINPWPHAKNRVGTGMYKYD--ENDDSGCRPIRYGVSCIGFDFMLKMDKYPFFNAFIGHKTPLESFADKIIEK-EEET-CEIG
                   *     :.:: .           *  ** *.*        ..   *  :  *.  *  .:    .  .:*:*::   : : :*:  : :    .  : * :
Prim.cons.         LL3SSV22VVA2P322PMI223PHL4NRPGKG4Y43DD344SD3TC4PL4YG4ACKS2D4QLD44K2PFFNSF442Y2LLEA442TII4K3E44TVC424

                          410       420       430       440       450       460       470       480       490       500
                            |         |         |         |         |         |         |         |         |         |
GP_RVFV            SA---HGNPCMKEKLVMKTHCPNDYQSAHYLNNDGKMASVKCPPKYGLTEDCNFCRQM---TGASL--KKGSYPLQDLFCQSSEDDGSKLKTKMKGVCEV
A0A143Q355_TOSV    PSTH-SSNACGREASLIKKKCGSNMSAFFYVNLAGQITVVKCDTNHVLSNDCGNCISK-TLSGQ-----KIYTPVQDMFCQKGWTESIPNTRYSKDLCSI
I1SV55_9VIRU       TTSPVTSESCLKSLRGIKHTCPPGFRSALFAHSTGKVRGIICDNQYQVTPDCKFCQST-ENNAE----VKNEIVIQDIVCQQNGTEYNGPIIKIPGYCGI
GP_PTPV            TN---KEFKCFEERAYIKGTCPTNINAVHYIDNKGKLRYVKCKENLEMTEDCAFCRKIKKKAGQSVQVQKTSVPLQDAICQENSDTYSGPKIPFKGVCKI
                   .        * ..   :*  *  .  :  :  .  *::  : *  :  :: **  * .     .      *    :** .**..            . * :
Prim.cons.         T422V4SN4C4KE444IK4TCP4N442AHY4NN4GK4R4VKCD4NY4LTEDC4FCRS4K334GQS2QV3K4S4PLQD4FCQ4N42EYSGPKIK4KGVC4I

                          510       520       530       540       550       560       570       580       590       600
                            |         |         |         |         |         |         |         |         |         |
GP_RVFV            GVQAHKKCDGQLSTAHEVVPFAVFKNSKKVYLDKLDLKTEENLLPDSFVCFEHKGQYKGTMDSGQTKRELKSFDISQCPKIGGHGSKKCTGDAAFCSAYE
A0A143Q355_TOSV    GLHTVKECKTG-TTNFERVGFIVVEGR-KMYIEQMKMRSRQEFSEDQFLCYKSE-SSSG------SSVKLKKVKVESCKGVTTSSASKCSGDEYFCSRYP
I1SV55_9VIRU       EDVKYRDCSDK-DVSFEKVAFALMKNK-KLYLPSMVLKYVDTVSADHFRCFRHK-KAHGVTGSNTDATDYVRVKPRECKQLDSSKTTKCTGDEVFCSYYN
GP_PTPV            GLIKYKECKFK-TSSYETVSFITLKEKGKIYIEHLMLKNIEVVTNVSFVCYEHVGQDEQEVE----HRALKRVSVNDCKIVDNSKQKICTGDHVFCEKYD
                        :.*.      .* * *  .:   *:*:  : ::  : .    * *:.   .                ..  .*  :     . *:**  **. *
Prim.cons.         GL4KYKECK4KLTTSFE4V4F2V4KNK2K4Y2E424LK44E4VS4DSFVC2EHKGQ44G333S2234R4LKRVKV44CK4VD4SK4KKCTGDEVFCS4Y4

                          610       620       630       640       650       660       670       680       690       700
                            |         |         |         |         |         |         |         |         |         |
GP_RVFV            CTAQYANAYCSHANGSGIVQIQVSGVWKKPLCVGYERVVVKRELSAKPIQRVEPCTTCITKCEPHGLVVRSTGFKISSAVACASGVCVTGSQSPSTEITL
A0A143Q355_TOSV    CETANVEAHCILRRHSAVIEVNVNGVWIVPRCIGYEEVLVRRTSLKVEDTSSRECDTCLWECGKNKLIVKTHGPKIVYATACSHGSCKSVMQKPATFVYL
I1SV55_9VIRU       CDKERSDAFCFHANGSGIIQVQYGGSWLQPSCVGFETVMVTKPRLSLPLSANLNCPSCVAKCHQNSLEINSNGFLITSAIACSHGECKTKTQKPRHSIIM
GP_PTPV            CSTSYPDVTCIHAPGSGPLYINLMGSWIKPQCVGYERVLVDREVKQPLLAPEQNCDTCVSECLDEGVHIKSTGFEITSAVACSHGSCISAHQEPSTSVIV
                   *     :. *     *. : ::  * *  * *:*:* *:* :            * :*: :*  . : :.: *  *  * **: * * :  *.*   : :
Prim.cons.         C4T4Y4DA4CIHANGSGIIQ22V4G2WIKP4CVGYERVLV4RE4L44PL4444NCDTCV42C44NGL42KSTGFKITSAVACSHGSCK244QKPSTS2IL

                          710       720       730       740       750       760       770       780       790       800
                            |         |         |         |         |         |         |         |         |         |
GP_RVFV            KYPGISQSSGGDIGVHMAHDDQSVSSKIVAHCPPQDPCLVHGCIVCAHGLINYQCHTALSAFVVVFVFSSIAIICLAVLYRVLKCLKIAP---RK----V
A0A143Q355_TOSV    PYPGNSEIVGGDIGVHMTEESSPSNIHMVAHCPAKDSCEVSSCLFCVHGLLNYQCHTLFS----ALLISTTVMSILTLLLLVKGAKDLV---KRLFYWL
I1SV55_9VIRU       QKPSSLKISGGEVGVHLSTEGEEPSYKLSIHCDPINQCDAYSCTFCWENVLNFHCHTLISSFVLALMIGSVASCALAIMVKISKGTKSVILKSRNPVLWV
GP_PTPV            PYPGLLASVGGRIGIHLSHTSDSASVHMVVVCPPRDSCAAHNCLLCYHGILNYQCHSTLSAILTSFLLILFIYTVFSVTTNILYVLRLIP---KQ----L
                     *.     ** :*:*::  ..  . ::   * . : * . .* .* ..::*::**: :*     :::       :::   :    :      :.    :
Prim.cons.         PYPG42422GGDIGVH2SHES4S4S42MVAHCPP4DSC42HSCLFC4HGLLNYQCHTLLSAFV3A2LISS4A444LAVL44ILKGLK44PLKS24222W2
```

```
                         810       820       830       840       850       860       870       880       890       900
                           |         |         |         |         |         |         |         |         |         |
GP_RVFV           LNPLMWITAFIRWIYKKMVARVAHNINQVNREIGWMEGGQLVLGN-----PAPIPRH-A----PIPRYSTYL-ML-LLIVSYASACSELIQASSRITTCS
A0A143Q355_TOSV   ITPLCWLSVFCGWVIKSWKKRVGSAISRTNDTIGWR--DNRRYRQ-----DVERAQYTGGA--PGAKYSFYG-VMVLGLLGNVHSCSESIIADSKIMQCT
I1SV55_9VIRU      IKLVKWLWL---QIWKALNKGMTTLSRKINDQTDLEVGSNQSFKNGIPLREVVVQKRTGGTRIKPVNYYLYGSTIVLGLITGSFSCTENLIASSKISSCF
GP_PTPV           KSPVGWLKLFINWLLTALRIKTRNVMRRINQRIGWVDHHDV-----------ERPRHRE----PMRRFKTTL-LLTLIMMTGGNACSNTVVANSKQTRCV
                   . : *:      : .           : *   .     :               :          .:      : * ::    :*:: : *.*:   *
Prim.cons.        I4P24WL4LFI3WI4KAL4KRV444IRRIND4IGW43G4N4333NGIPLR3VERPRHTGG2RIP44RYSTY2S4LVLGL4TG442CSE4IIASSKIT4C4

                         910       920       930       940       950       960       970       980       990      1000
                           |         |         |         |         |         |         |         |         |         |
GP_RVFV           TEGVNTKCRLSGTALIRAGSVGAEACLMLKGVKEDQTKFLKIKTVSSELSCREGQSYWTGSISPKCLSSRRCHLVGECHVNRCLSWRDNETSAEFSFVGE
A0A143Q355_TOSV   TSGSSTLCKASGTVIMKLGPIGSESCLILKGLKDSEKQFISIKTISSELTCREGESFWTTLYTPVCLSSRRCHLMGECVSDRCLKWKTNETSAEFTGKAH
I1SV55_9VIRU      IESGRHVCKLSGVVNLRVGTIGSESCLIVKGPLDGQTQALKIRTKSSELVCQEGQSFWTNHFTPKCYSSRRCHLVGECQASNCLEWNASTVSQEFRSQYE
GP_PTPV           QEGSNTKCSITATITLRAGVIGAESCFIIKGPMENQQKTISIKTISSETVCREGSSFWTSLYIPSCLSSRRCHLVGDCVGNKCQSWRDDQLSREFSGVKD
                   ..    *  :..  :: * :*:*:*:::**  :.: : :.*:* ***  *:**.*:**    * * *******:*:*  ..* .*. .  * **    .
Prim.cons.        TEGSNTKCKLSGTV4LRAG4IG2ESCLILKGPK24QT2F22IKTISSELVCREGQSFWT4LYTPKCLSSRRCHLVGECV4NRCLSWRDNETSAEFSGV4E

                        1010      1020      1030      1040      1050      1060      1070      1080      1090      1100
                           |         |         |         |         |         |         |         |         |         |
GP_RVFV           STTMRENKCFEQCGGWGCGCFNVNPSCLFVHTYLQSVRKEALRVFNCIDWVHKLTLEITDFDGSVSTIDLGASSSRFTNWGSVSLSLDAEGISGSNSFSF
A0A143Q355_TOSV   GEVVHENRCFEQSGGIGYGCFNVNPSCLYVHSYLKSVYKNGFKIFRCVAWNHRIRLEVTTHSRKFPMV-LMAMSTQPTDWGSIGLILDSEGITGTNAYSF
I1SV55_9VIRU      PGMIVENVCFEQCGGIGCSCFNVNPSCLFGMTILKPTYDKAIKVFQCASWTHRITLEFEGPGITAKQVSLTSMSTQVAEWGSITLSVDADSTIDNPNISF
GP_PTPV           NHIMNENKCFEQCGAIGCGCFNINPSCLYVHAYLKSARNEAVRVFSCSDWVHRVSFEVKGPDGETELVTLGSPGTKFLNWGTLSLSLDAEGISGTNSISF
                     : ** ****.*. * .***:*****:  : *:.. .:..::* *  * *:: :*.   .     : * : .::  :**:: * :*::.  ..   **
Prim.cons.        444M4ENKCFEQCGGIGCGCFNVNPSCL2VHTYLKSV2KEA42VF4C4DWVHRITLEVTGPDG4444V3LG2MSTQFTNWGSISLSLDAEGISGTNSISF

                        1110      1120      1130      1140      1150      1160      1170      1180      1190      1200
                           |         |         |         |         |         |         |         |         |         |
GP_RVFV           IESPSKGYAIVDEPFSEIPRQGFLGEIRCNSESSVLSAHESCLRAPNLISYKPMIDQLECTTNLIDPFVVFERGSLPQTRNDKTFAASKGNRGVQAFSKG
A0A143Q355_TOSV   MKHGSGSFAIIDDPYSPEPRKGFLGEVRCPTEETAIRASPSCKMAPNLIEYQPEMDTAECTTNMIDPMAIFNRGSLPQVRDGMTFTQSIEKNTVQALTTG
I1SV55_9VIRU      LKTREGEYALIEESITTSPQKGRLGEIRCQTEQQAAIGSPSCLQADKLVDYRPQYDTLECISKLIDPRAILKRNGLPQSRGKYLYTPSLGSDTVQAVSRN
GP_PTPV           LESSKGGFALYDEGYNEIPREGFLGEIRCSSESAAISAHKSCIRAPGLIKYKPMTDQIECTASLVDPFAIFLKGSLPQTRNGQTFTSTKDKKTVQAFTNG
                  ::  .  :*: ::  .  *::* ***:** :*. .  .  **  *  *:.*:*  *  ** :.::** .:: :..*** *.   :: :  .  ***.: .
Prim.cons.        L2S4SGG2A2IDEPYSEIPRKGFLGEIRC42ES4AISA2PSCLRAPNLI4YKPM4D2LECTTNLIDPFAIF4RGSLPQTRNG4TFT4SKGK4TVQAF24G

                        1210      1220      1230      1240      1250      1260      1270      1280      1290      1300
                           |         |         |         |         |         |         |         |         |         |
GP_RVFV           SVQADLTLMFDNFEVDFVGAAVSCDAAFLNLTGCYSCNAGARVCLSITSTGTGSLSAHNKDGSLHIVLPSENGTKDQCQILHFTVPEVEEEFMYSCDGDE
A0A143Q355_TOSV   EVRASIRLTLDDYDVVYQNSQSDCSATFKNLTGCYSCDEGSRMCYQVKAEGETIFHFINEDESINVINKVNPDVSDYCTVLHFSRPVISIEGRYDCGLSK
I1SV55_9VIRU      TIEATMTLTLDNYEVSFLSEKTECLASFVNISGCYSCHSGASICFKVTSKTPASIYIRDEKETKIISKSITAGEKVICSVLHYSKPIVDEIFEYTCGSEW
GP_PTPV           AIKALLSINLDDHEIVFINKVKNCDATFLNVSGCYSCDYGAHVCVKVKSSESADFFAESEDKTTVLSFPIQSGTHDYCQVLHFQKPLVDERLSYSCGSEP
                   :.* : : :*:.:: : .   .* *:* *::*****. *: :* .:.:     :   .:. :  :      .    * :**:  * :.    * *. .
Prim.cons.        424A4LTLTLD2YEVVF4N4444CDATFLNL2GCYSCD4GARVC4KV2S4G4ASF4A4NEDE244IS4PI44GTKDYCQVLHFSKP4VDEEF4YSCGSE4

                        1310      1320      1330      1340      1350      1360      1370      1380      1390      1400
                           |         |         |         |         |         |         |         |         |         |
GP_RVFV           RPLLVKGTLIAIDPFDDRREAGGESTVVNPKSGSWNFFDWFSGLMSWFGGPLKLYSSFACMLHYQLGSFSSLYILEEQASLKCGLLPLRRPHRSVRVKVI
A0A143Q355_TOSV   RPMVIKGTLIATAPHDDRKHEGGSSLIVNPKGGTVDFFGWVSGLTSWLGGPLKTFLTILGFLALGLLLVVIIIII------------ARTGVQQALKKKN
I1SV55_9VIRU      KKIQVHGSLILLSLIDDHK-EGGSTIVVGPSSSSFSVGDWTTGLLSWFGGPIKTLTLLIIYVVATIIVIALVVAL------------IKLALKTMILKKN
GP_PTPV           KLIVIKGTLVCMGVYDFRNKTGGSSTVVNPSEGAWSISNWFSGLLDWLGGPMKAILKILGFIAIGIVCFVLFMIL------------IRIAVNSINIKKK
                  : : ::*:*:     * :.  **.: :*.*. .: .. .* :** .*:***:*    :   :   :  .  .  :             :   .    *
Prim.cons.        2PIV2KGTLIA44P4DDRK3EGGSSTVVNP2SGSWSFFDWFSGLLSW2GGPLKT4L4ILGFLA4G244FVL44ILEEQASLKCGLLPIR4AV4S444KKN


GP_RVFV           C
A0A143Q355_TOSV   K
I1SV55_9VIRU      V
GP_PTPV           N

Prim.cons.        4
```

**Table S6: Sequence alignment of *Phenuiviridae* NSm**. Red type/ asterisk (*), green type/colon (:), and blue type/dot (.) indicate identical amino acid residues, conserved substitution, and semi-conserved substitutions respectively.

```
                          10        20        30        40        50        60        70        80        90       100
                           |         |         |         |         |         |         |         |         |         |
GP_RVFV           MY--VLLTILISVLVCEAVIRVSLSS-----TREETCFGDSTNPEMIEGAWDSLREE---EMPEELSCSISGIREVKTSSQELYRALKAIIAAD--GLNN
A0A143Q355_TOSV   MF--IAKLLLISCLVQQAYLKYTIISEIGNGTRTSFCYSSRTSLGSLLEAWDNITES---IGLTSGTCKV-GSLENRDCKRDIMSNYVAEVHSNLHGLGA
I1SV55_9VIRU      ------------------MHQITVVSG---PTEVSTCFGSLHPFQSLKPVMANALGV----LEGKMFCSI-GGRSLEEDDEGVLAVVQGLQDSQ--QEEE
GP_PTPV           MIFTILNVLTRAMLVMSMYSLTTWDST----SRNDMCFSNDSPLEGLVYYWETHSKRHDYKKQESQRCRV-GDSDKKMITNVTIISLISEIQKSISELSL
                                        :  *     :. . *:..      :     .           .  * : *  . .   .       .    .
Prim.cons.        M3FTIL33LLIS3LV33AY444T44S3IGN2TR4STCF2S4TPLESL44AWDN44E4HDY344ES44CS2SG4RE4K444444444L4AEI4S422GL44

                         110       120       130       140       150       160       170       180
                           |         |         |         |         |         |         |         |
GP_RVFV           ITCHGKDPEDKISLIKGP-PHKKRVGIVRCERRRDAKQIGRETMAGIAMTVLPALAVFALAPVVFA-----------------
A0A143Q355_TOSV   MM-HNYGDDKRVEVLNDVIPDELNPGRHNCS-DLDRKPFWTEFID--------------------------------------
I1SV55_9VIRU      ISCHGTPKGKLVKYKRHPRKASGKKGAVDCVSGKELKPQKKRGDLAHISEHFQADEVRSSSVNISTTLKTILVAALLTSALVN
GP_PTPV           SCVNDDNSTGQVLTFNGLEDTIRGDYIVDCVTGLYQSDIGVGVGLGRTHHGH-------------------------------
                     :.      :   .             *      .
Prim.cons.        I4CHG4444K4V444NGP3P44444GIVDCV3GLD4KPIG4E44LG3333332A22V222222222TLKTILVAALLTSALVN
```

**Table S7: Sequence alignment of *Phenuiviridae* Gn$^{cyto}$.** Red type/ asterisk (*), green type/colon (:), and blue type/dot (.) indicate identical amino acid residues, conserved substitution, and semi-conserved substitutions respectively.

```
                           10        20        30        40        50        60        70        80        90       100
                            |         |         |         |         |         |         |         |         |         |
GP_RVFV           AVLYRVLKCLKIAPRKVLNPLMWITAFIRWIYKKMVARVAHNINQVNREIGWMEGGQLVLGNPAPIPRH-APIPRY-------STYLMLLLIVSYASA--
A0A143Q355_TOSV   LLVKGAKDLVKRLFYWLITPLCWLSVFCGWVIKSWKKRVGSAISRTNDTIGWRDNRRYRQDVERAQYTGGAPGAKY-------SFYGVMVLGLLGNVHS-
I1SV55_9VIRU      AIMVKISKGTKSVILKSRNPVLWVIKLVKWLWLQIWKALNKGMTTLSRKIN--DQTDLEVGSN-QSFKNGIPLREVVVQKRTGGTRIKPVNYYLYGSTIV
GP_PTPV           SVTTNILYVLRLIPKQLKSPVGWLKLFINWLLTALRIKTRNVMRRINQRIGWVDHHDV------ERPRHREPMRRF-------KTTLLLTLIMMTGGN--
                   :        :       .*: *:  :  *:           :   .  *.  :                 *  .
Prim.cons.        AV444ILK4LK44P4KL4NP24WL44FI4WL4K444KRV44424R4NR4IGW3D44DL33G33244PRHGAP4RRYVVQKRTGSTYL4LVLI4LYGS42V


GP_RVFV           ----
A0A143Q355_TOSV   ----
I1SV55_9VIRU      LGLI
GP_PTPV           ----

Prim.cons.        LGLI
```

**Table S8: Sequence alignment of RVFV Gn$^{cyto}$ and NSm.** Red type/ asterisk (*), green type/colon (:), and blue type/dot (.) indicate identical amino acid residues, conserved substitution, and semi-conserved substitutions respectively.

```
                       10        20        30        40        50        60        70        80        90       100
                        |         |         |         |         |         |         |         |         |         |
GP_RVFV_Gncyto-------------------------------------AVLYRVLKCLK--IAPRKV---LNPLMWITAFIRWIYKKMVARVAHN-INQV-------NREI
GP_RVFV_NSm   MYVLLTILISVLVCEAVIRVSLSSTREETCFGDSTNPEMIEGAWDSLREEEMPEELSCSISGIREVKTSSQELYRALKAIIAADGLNNITCHGKDPEDKI
                                                    ::  . ..*:    *.::   :. :  :.:  : :*: : * :* : :*::       : :*
Prim.cons.    MYVLLTILISVLVCEAVIRVSLSSTREETCFGDSTNP222222222L2EE22P222SCS22222222222222Y2222A22A22G2N22TCHGKDP222I

                      110       120       130       140       150
                        |         |         |         |         |
GP_RVFV       GWMEG--GQLVLG--------NPAPIPRHAPIPRYSTYLMLLLIVSYASA---
GP_RVFV       SLIKGPPHKKRVGIVRCERRRDAKQIGRETMAGIAMTVLPALAVFALAPVVFA
              . ::*   :  :*        :.  * *.:      * *  *  :.: *..
Prim.cons.    2222GPP22222GIVRCERRR2222I2R22222222T2L22L22222A22VFA
```
